# microRNA-138 controls hippocampal interneuron function and short-term memory

**DOI:** 10.1101/2021.05.12.443573

**Authors:** R. Daswani, C. Gilardi, M. Soutschek, P. Nanda, K. Weiss, S. Bicker, R. Fiore, C. Dieterich, P.L. Germain, J Winterer, G Schratt

## Abstract

The proper development and function of neuronal circuits relies on a tightly regulated balance between excitatory and inhibitory (E/I) synaptic transmission, and disrupting this balance can cause neurodevelopmental disorders, e.g. schizophrenia. microRNA-dependent gene regulation in pyramidal neurons is important for excitatory synaptic function and cognition, but its role in inhibitory interneurons is poorly understood. Here, we identify miR-138-5p as a regulator of short-term memory and inhibitory synaptic transmission in the hippocampus. Sponge-mediated miR-138-5p inactivation specifically in parvalbumin (PV)-expressing interneurons impairs spatial recognition memory and enhances GABAergic synaptic input onto pyramidal neurons. Cellular and behavioural phenotypes associated with miR-138-5p inactivation are paralleled by an upregulation of the schizophrenia-associated *Erbb4*, which we validated as a direct miR-138-5p target gene. Our findings suggest that miR-138-5p is a critical regulator of PV interneuron function, with implications for cognition and schizophrenia. More generally, they provide evidence that microRNAs orchestrate neural circuit development by fine-tuning both excitatory and inhibitory synaptic transmission.

## Introduction

microRNAs (miRNAs) are short non-coding RNAs which act as negative regulators of mRNA translation and stability (Bartel, 2018). Over the last decade, a large body of evidence shows that miRNAs control excitatory neuron development, function and plasticity (McNeill and Van Vactor, 2012; Schratt, 2009). Specific miRNAs, e.g. miR-132, -134 and -138, have been identified which control dendritic spines, the major sites of excitatory synaptic contact (Schratt et al., 2006; Siegel et al., 2009). Complete lack of miRNAs from excitatory forebrain neurons in Dicer-deficient mice enhances learning and memory (Konopka et al., 2010), and specific miRNAs have been linked to both short-term and long-term memory regulation (Cheng et al., 2018; Gao et al., 2010; Walgrave et al., 2021). Although miRNAs are likewise substantially expressed in inhibitory γ-aminobutyric acid (GABA)ergic interneurons (He et al., 2012), relatively little is known about miRNA function in this cell-type. The complete lack of miRNAs reduces the number of cortical interneurons (Tuncdemir et al., 2015) while the absence of miRNAs in interneurons expressing vasoactive intestinal peptide (VIP) leads to cortical circuit dysfunction (Qiu et al., 2020). miR-128 knockout in Drd1a-positive GABAergic medium spiny neurons of the striatum results in increased motor activity and fatal epilepsy (Tan et al., 2013).

In the rodent hippocampus, microcircuits of excitatory pyramidal neurons and local inhibitory interneurons provide an extensively studied model in the context of information processing, with specific implications for the control of spatial short-term and long-term memory (Booker and Vida, 2018; Markram et al., 2004; Pelkey et al., 2017). Among the different interneuron classes, fast-spiking parvalbumin (PV) expressing interneurons play a particularly prominent role in controlling pyramidal neuron output to drive appropriate behavioral responses (Murray et al., 2011; Rico and Marin, 2011). For example, CA1 PV interneurons are required for spatial working memory, but neither for reference memory nor for memory acquisition in contextual fear conditioning (CFC) (Fuchs et al., 2007; Lovett-Barron et al., 2014; Murray et al., 2011). PV interneuron dysfunction has been implicated in neuropsychiatric disorders, most notably schizophrenia, but also epilepsy and autism-spectrum disorders (ASD) (Del Pino et al., 2018; Sohal and Rubenstein, 2019). However, the role of specific miRNAs in PV interneurons in the context of higher cognitive function is completely elusive.

## Results

We previously identified the brain-enriched miR-138-5p as an important regulator of excitatory synapse function in hippocampal pyramidal neurons (Siegel et al., 2009). To study the role of miR-138-5p on a behavioral level, we generated mice with a conditional ROSA26 transgene (138-floxed) which allows expression of a miR-138-5p inactivating sponge transcript harboring a lacZ coding sequence and 6 imperfect miR-138-5p binding sites (6x-miR-138sponge) upon Cre-recombinase expression. Sponge transcripts sequester endogenous miRNA, thereby leading to miRNA inactivation and the de-repression of cognate target genes (Ebert and Sharp, 2010). 138-floxed mice without Cre transgene served as a control throughout the experiments. Since these control mice completely lack expression of any sponge-related transcript *in vivo*, we first carefully validated specificity and efficiency of the 138-sponge transcript in primary rat hippocampal neurons. Therefore, we compared the 138-sponge transcript to an identical control sponge, except that the 6 imperfect miR-138 binding sites were replaced with a shuffled sequence not supposed to sequester any miRNA expressed in neurons (cf. Methods). This analysis revealed a robust and specific inactivation of miR-138-5p by 138-sponge *in vitro* (**suppl. Fig. 1a, b, c**).

We then activated 138-sponge expression at embryonic stage *in vivo* by crossing 138-floxed mice to the ubiquitous Cre-driver line CMV-Cre (138-sponge^ub^, **Fig. 1a**). lacZ staining revealed highly penetrant expression of 6x-miR-138 sponge in the hippocampus of 138-floxed mice upon CMV-Cre expression (**suppl. Fig. 1d**). To test for the degree of miR-138-5p inhibition *in vivo* we made use of a dual fluorescence miR-138 sensor virus which we injected into the hippocampus of 138-sponge^ub^ mice. Briefly, the sensor consists of a GFP coding sequence (cds) followed by two perfect miR-138 binding sites and a mCherry cds (suppl. Fig. 1e). Thus, the lower the ratio of GFP/mCherry, the higher the cellular miR-138 activity. Quantifying GFP/mCherry ratios over many infected neurons revealed that in neurons with 6x-miR-138-sponge expression, GFP/mCherry ratios were significantly higher compared to controls, indicative of an efficient sequestering of endogenous miR-138-5p by our sponge construct (**Fig. 1b**).

**Figure 1.**
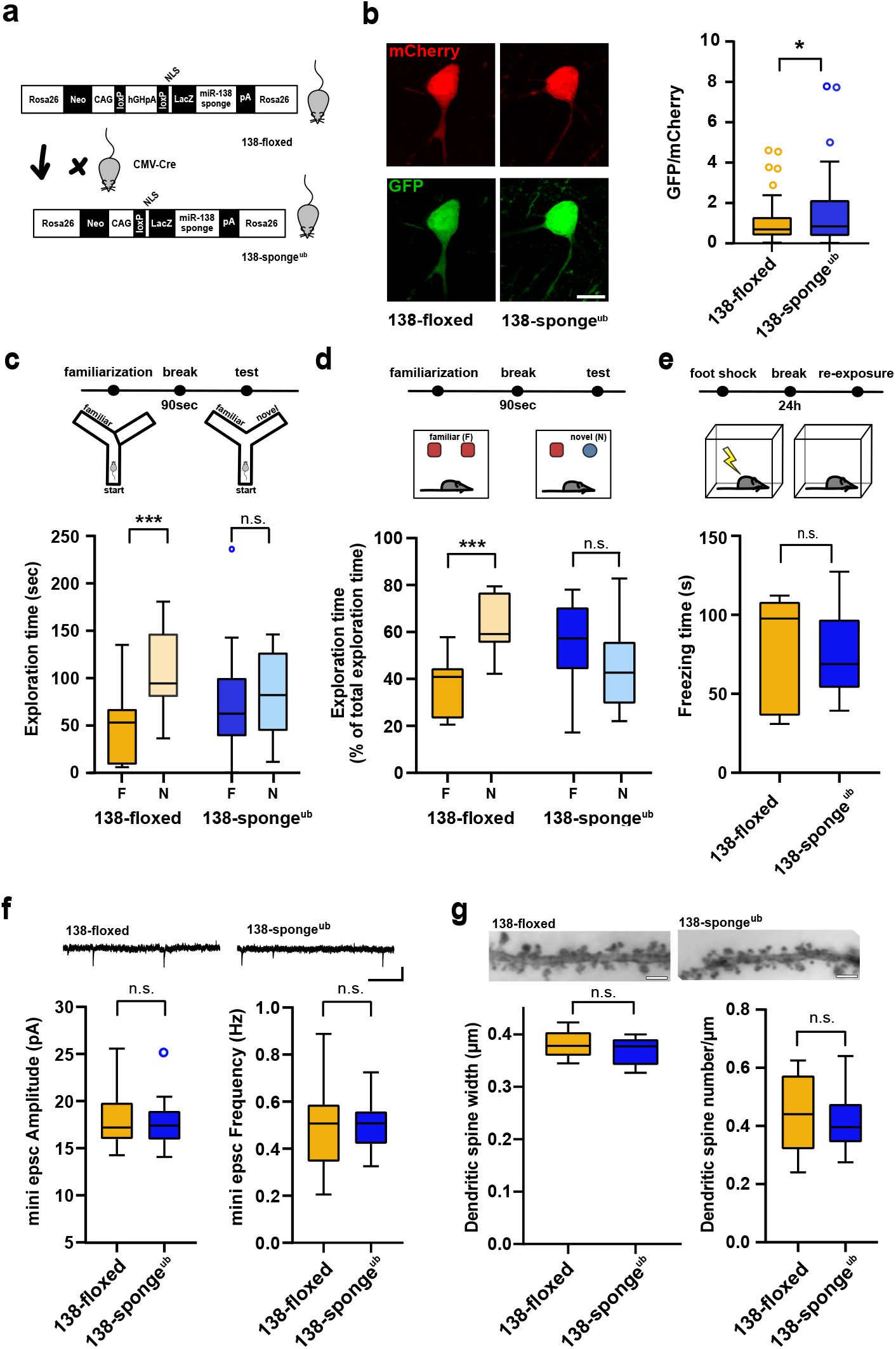
**(a)** Schematic overview of the strategy for generating 138-sponge^ub^ mice. **(b)** Left: representative images of mCherry and GFP expression in hippocampal neurons from 138-floxed and 138-sponge^ub^ mice, respectively. Right: bar graphs of GFP/mCherry ratios from CA1 hippocampal neurons infected with a 138-pbds sensor construct; 138-floxed: n=105 cells from two mice, 138-sponge^ub^: n=127cells from three mice; p= 0.02 (KS-test). **(c)** Upper panel: schematic representation of the Y maze novelty preference task; lower panel: exploration time spent in familiar (F) and novel (N) arm; 138-floxed n=12 mice; 138-sponge^ub^ n=14 mice; ***p=0.005; n.s.=0.711 (Student’s two-tailed heteroscedastic t test). **(d)** Upper panel: schematic representation of the novel object recognition task; lower panel: exploration time presented as percentage of total time spent with either novel or familiar object; 138-floxed n=12 mice; 138-sponge^ub^ n=12 mice; **** p<0.00002; n.s. p=0.11 (Student’s two tailed heteroscedastic t test). **(e)** Upper: schematic representation of the contextual fear conditioning task; lower: time (s) mice spent freezing 24 h after the foot shock was administrated; 138-floxed n=7 mice; 138-sponge^ub^ n=7 mice; n.s. p=0.97 (Student’s two-tailed heteroscedastic t test). **(f)** mEPSC recording in CA1 pyramidal neurons. Upper panel: example traces; scale bar: 20 pA, 500 ms. Lower panel left: mEPSC amplitude (138-floxed: range, from 14.3 to 25.6 pA; median, 17.2 pA; interquartile range [IQR], 3.9 pA. 138-sponge^ub^: range, from 14.1 to 25.2 pA; median, 17.4 pA; IQR, 3.1 pA; n.s. p=0.74 Student’s two-tailed heteroscedastic t test). Lower panel right: mEPSC frequency (138-floxed: range, from 0.2 to 0.9 Hz; median, 0.5 Hz; IQR, 0.2 Hz. 138-sponge^ub^: range, from 0.3 to 0.7 Hz; median, 0.5 Hz; IQR, 0.1 Hz; n.s. p=0.91 Student’s two-tailed heteroscedastic t test). 138-floxed n=13 cells/4mice; 138-sponge^ub^ n=13cells/3mice. **(g)** Upper panel: representative images of Golgi-stained CA1 pyramidal neuron dendritic segments of the indicated genotypes. Lower panel: quantification of dendritic spine width (left) and density (number/μm; right) based on Golgi staining; 138-floxed: n=15 cells from 3 mice (1312 spines total); 138-sponge^ub^ n=18 cells from 3 mice (1687 spines total) (n.s., p=0.25 (width); p=0.49 (density); Student’s two tailed heteroscedastic t test). **Figure 1 – source data:** this file contains the raw data on which the graphs in Fig. 1 are based.

We next assessed cognitive abilities of 138-sponge^ub^ mice using behavioral testing. Locomotion in the home cage was similar between 138-sponge^ub^ and control mice, ruling out severe developmental motor impairments as a potential confound (**suppl. Fig.1f**). In the Y-maze test, no genotype-dependent differences in spontaneous alternations were observed, suggesting that exploratory behavior was not affected by miR138-5p inactivation (**suppl. Fig. 1g**). In contrast, 138-sponge^ub^ mice displayed a significant impairment in novelty preference (**Fig. 1c**), indicating a loss of spatial short-term memory. In the novel object recognition (NOR) task, 138-sponge^ub^ mice were unable to discriminate the novel from the familiar object (**Fig. 1d**), thereby corroborating the observed short-term memory deficit. In contrast, associative long-term memory, as assessed by classical fear conditioning (**Fig. 1e**), as well as anxiety-related behavior (open field, EPM), was not affected by miR-138-5p inhibition (**suppl. Fig. 1h, I, j**). Thus, ubiquitous miR-138-5p inhibition leads to a short-term memory deficit.

We went on to test whether short-term memory impairments in 138-sponge^ub^ mice were associated with alterations in synaptic transmission in hippocampal area CA1 and recorded miniature excitatory postsynaptic currents (mEPSCs) in CA1 pyramidal neurons. Amplitude and frequency of mEPSCs were indistinguishable between 138-sponge^ub^ and control slices (**Fig. 1f**, **suppl. Fig. 1k, 1l**). Likewise, we did not detect any significant alterations in dendritic spine morphology in these neurons using Golgi staining (**Fig. 1g**). Finally, we did not observe differences in paired-pulse ratio (PPR), which negatively correlates with presynaptic release probability (**suppl. Fig. 1m**), suggesting that excitatory synaptic transmission at the Schaffer collateral CA1 pyramidal cell synapse was not affected by miR-138-5p inhibition.

To identify miR-138-5p target mRNAs and to obtain further insight into the biological function of miR-138-5p regulated genes, we performed polyA-RNA sequencing with total RNA isolated from hippocampal tissue. Differential gene expression analysis recovered a total of 338 differentially expressed genes (DEG; 265 upregulated, 73 downregulated) (FDR<0.05; **Fig. 2a**). The presence of miR-138-5p binding sites correlated with increased transcript levels compared with 138-floxed mice (**Fig. 2b**). miR-138-5p 7-mer 1a, and to a lesser extent 7mer-m8 and 8mer sites, predicted significant de-repression (**suppl. Fig. 2a**). In total, 56 (21%) of the upregulated, but only 6 (8%) of the downregulated genes harbor miR-138-5p binding sites within their 3’UTR. Together, this data demonstrates that many of the observed upregulated genes are a direct consequence of miR-138-5p inhibition and further confirms the specificity of the miR-138-5p sponge transgene.

**Figure 2.**
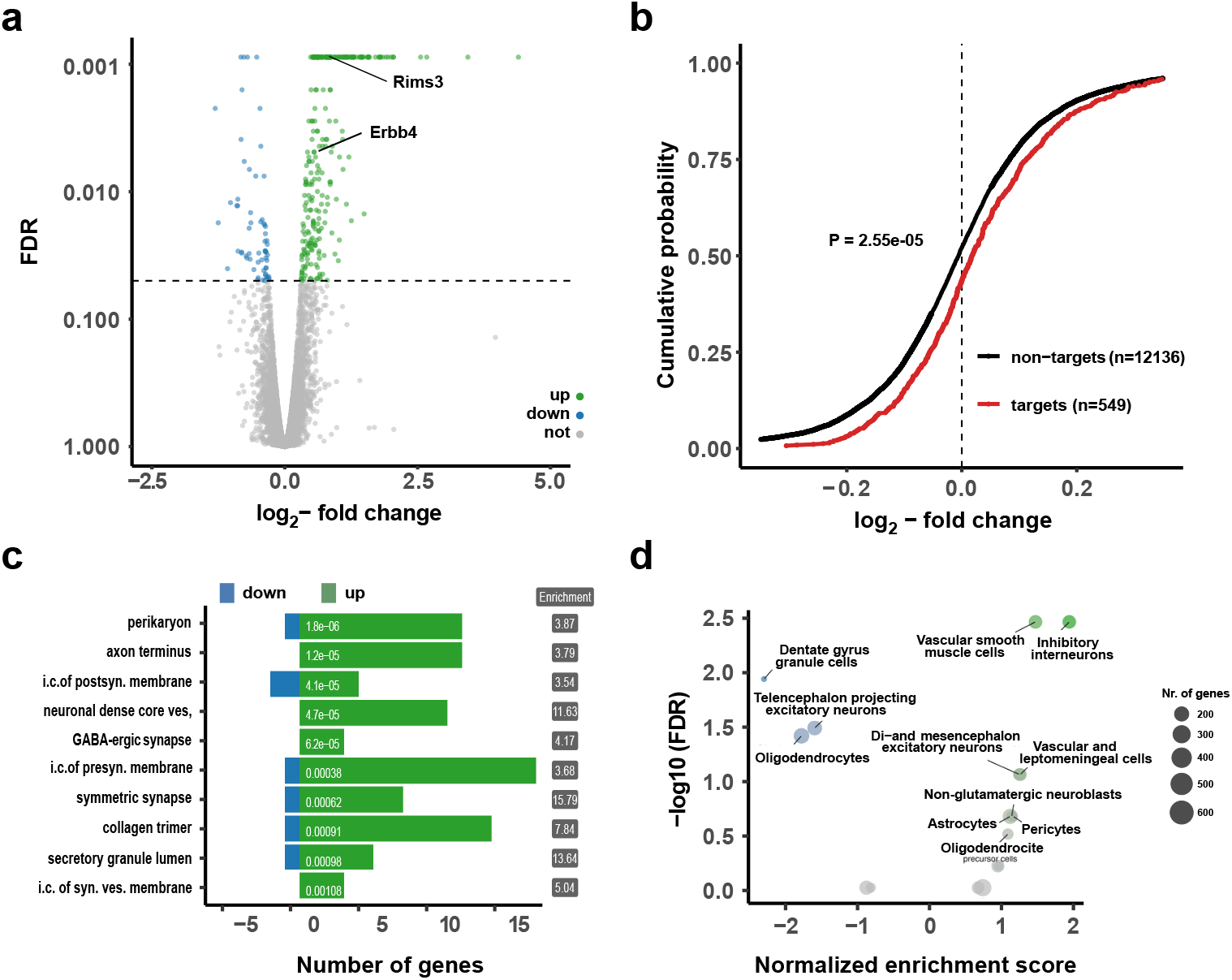
**(a)** Volcano plot of differentially expressed genes (DEGs) obtained from polyA-RNAseq of total hippocampal RNA from 138-flox and 138-sponge^ub^ mice. N=3. Genes with FDR<0.05 are labeled blue (downregulated) or green (upregulated). Rims3 and Erbb4 are indicated. **(b)** Cumulative distribution plots of log_2_-fold expression changes (138-sponge^ub^/138-floxed) for genes either containing (targets, red curve) or not containing (non-targets, black curve) predicted miR-138 binding sites. p=2.55e^-05^ (KS-test). **(c)** Gene ontology (GO) term analysis for DEGs. Top ten enriched cellular component (CC) GO terms with less than 200 total genes are shown. **(d)** Enrichment analysis of DEGs in different brain cell types based on published single-cell RNA-seq data(Zeisel et al., 2018). Normalized enrichment score>0: upregulated in 138-sponge^ub^ mice.

Gene ontology (GO) term analysis on DEGs from the RNA-seq analysis (**Fig. 2c**; **suppl. Fig. 2b**) revealed that many GO terms associated with synaptic function are strongly over-represented in genes upregulated in the hippocampus of 138-sponge^ub^ mice. In order to specify the origin for the observed gene expression changes, we compared DEGs to single cell RNA-seq data from different cell types present in the hippocampus (Zeisel et al., 2018) (**Fig. 2d, suppl. Fig. 2c**). Surprisingly, we found that those genes which were significantly upregulated in 138-sponge^ub^ mice are strongly enriched in inhibitory GABAergic interneurons. This finding is in line with the observation that excitatory synaptic transmission in hippocampal CA1 was not affected by miR-138-5p inactivation (**Fig. 1f, g**, **suppl. Fig. 1 j-l**). Accordingly, many of the upregulated miR-138-5p targets showed a strong expression signal in different classes of inhibitory interneurons (**suppl. Fig. 2d**). In contrast, known validated targets of miR-138-5p which are not enriched in inhibitory interneurons (e.g., Lypla1, Sirt1, Fmr1) were not differentially expressed between 138-sponge^ub^ and control mice (**suppl. Fig. 2e**).

Next, we performed single-molecule miRNA FISH in cultured rat hippocampal neurons to visualize miR-138-5p expression at subcellular resolution. This analysis revealed strong expression of miR-138-5p in both Camk2a-positive excitatory and Erbb4-positive inhibitory hippocampal neurons (**Fig. 3a**). Erbb4-positive cells express Gad65, but not Camk2a, confirming their GABAergic phenotype (**Fig. 3a**). Quantification of the FISH signal didn’t reveal any significant differences in the average cellular miR-138-5p expression between excitatory and inhibitory neurons (**suppl. Fig. 3a**), in agreement with an analysis of published miRNA-seq data from the mouse cortex (**suppl. Fig. 3b**) (He et al., 2012). In contrast, miR-138-5p expression was undetectable in GFAP-positive glial cells (**suppl. Fig. 3c**). Thus, miR-138-5p is robustly expressed in inhibitory interneurons, consistent with a previously unrecognized function in these cells.

**Figure 3.**
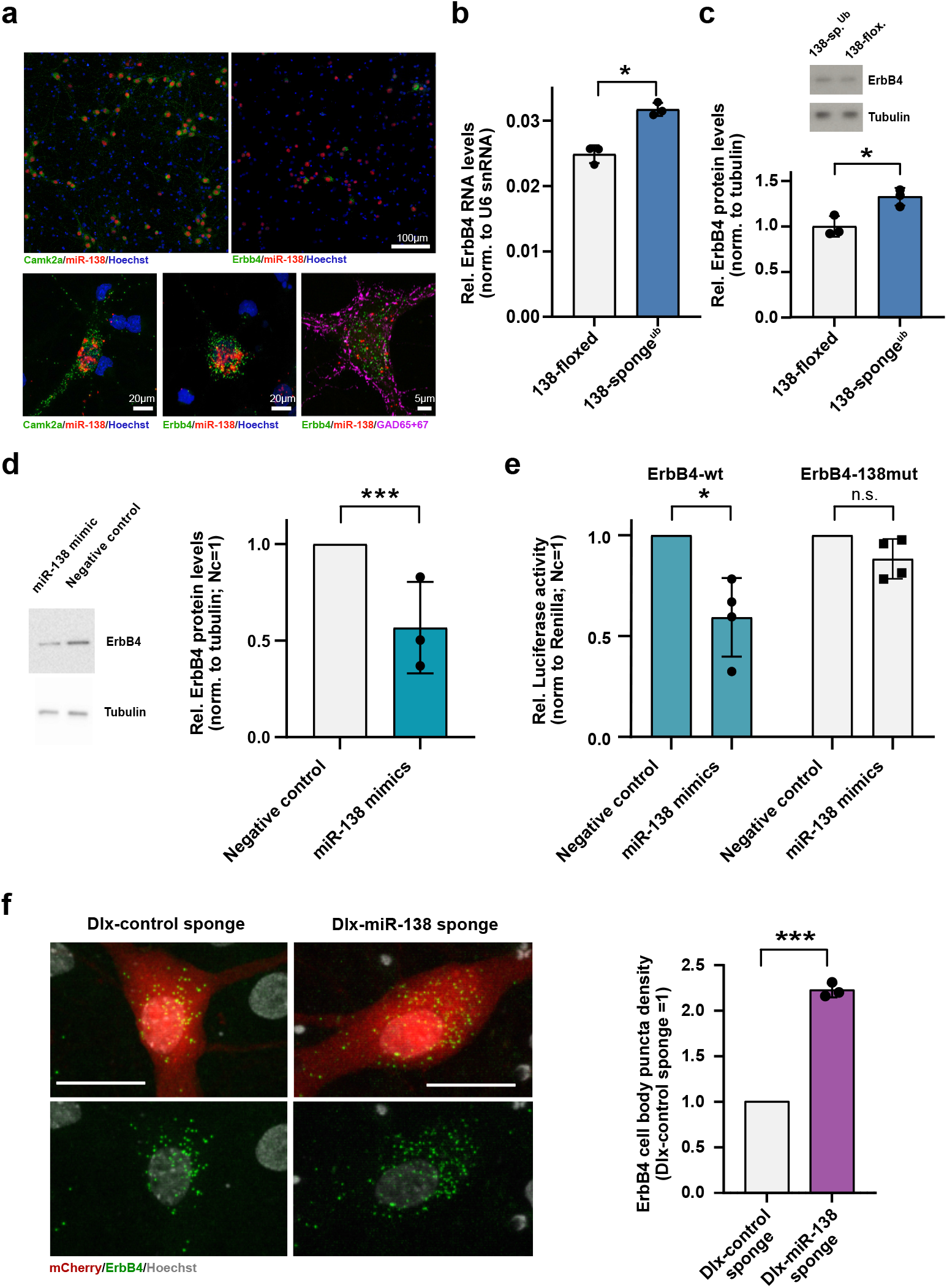
**(a)** Single-molecule (Sm) FISH analysis of miR-138 (red) together with Camk2a or Erbb4 mRNA to label excitatory or inhibitory neurons, respectively. Hoechst was used to counterstain nuclei. GAD65/67 antibody staining was used to identify GABAergic neurons. Scale bar=100μm (upper); 20μm (lower left and center), 5μm (lower right). **(b)** qPCR analysis of ErbB4 mRNA in total hippocampal RNA obtained from 138-floxed or 138-sponge^ub^ mice. U6 snRNA was used for normalization. n=3 mice; *p=0.003, (Student’s two-tailed heteroscedastic t test) **(c, d)** Western blot analysis of Erbb4 protein in hippocampal lysates from 138-floxed or 138-sponge^ub^ mice (c) or lysates from rat hippocampal neurons (DIV12) transfected with miR-138 or control mimic (d). Tubulin was used for normalization. (c) n=3 mice; *p=0.025 (student’s two-tailed heteroscedastic t test); (d) n=3 independent transfections; ***p=0.0009 (one sample t-test). **(e)** Relative luciferase activity in rat cortical neurons (DIV9-12) transfected with Erbb4 3’UTR constructs with (138mut) or without (wt) a mutation in the miR-138 binding site, together with miR-138 or negative control mimics. Negative control mimic = 1. n=4 independent transfections, *p=0.025, n.s. p= 0.09 (student’s two-tailed heteroscedastic t test). **(f)** Sm FISH analysis of Erbb4 (green) in rat hippocampal interneurons infected with Dlx-control-sponge or miR-138-sponge. Left panel: representative neurons, scale bar=20 μm. Green: Erbb4 FISH; grey: DAPI (nuclei); red: mCherry (Dlx5/6 expressing interneurons). Right panel: Erbb4 FISH quantification. Control sponge=1. N=3 independent infections (10-12 cells per condition), ***p=0.0004 (one-sample t-test). **Figure 3 – source data:** this file contains the raw data on which the graphs in Fig. 3 are based.

We went on to validate miR-138-5p dependent regulation of predicted interneuron-enriched target genes, focusing on Erbb4 (Erb-B2 receptor tyrosine kinase 4). Erbb4 is a SCZ risk gene which has been linked to the regulation of GABAergic synaptic transmission and short-term memory (Fazzari et al., 2010; Wang et al., 2018). Using qPCR and Western blot, we observed a significant upregulation of both Erbb4 mRNA **(Fig. 3b)** and protein **(Fig. 3c)** levels in the hippocampus of 138-sponge^ub^ compared to 138-floxed mice, thereby validating our results from RNA-seq. On the other hand, transfection of primary rat hippocampal neurons with a synthetic miR-138-5p mimic reduced Erbb4 protein levels **(Fig. 3d)**, demonstrating that miR-138-5p is necessary and sufficient for the inhibition of Erbb4 expression. In luciferase reporter gene assays, transfection of miR-138-5p significantly reduced the expression of an Erbb4 3’UTR construct containing a wild-type, but not mutant miR-138-5p binding site (**Fig. 3e**), rendering Erbb4 as direct miR-138-5p target. Furthermore, we were able to validate the presynaptic vesicle associated Rims3 as an interneuron-enriched miR-138-5p target **(suppl. Fig. 3d-f)**.

To study the regulation of Erbb4 by miR-138-5p specifically in interneurons, we designed a viral construct in which expression of an inhibitory miR-138 sponge or a respective control sponge is under the control of the interneuron-specific Dlx5/6 promoter (Dlx-138-sponge, Dlx-control sponge). rAAV expressing Dlx-138- or control sponge co-localizes with Gad65/67 in primary rat hippocampal neurons, confirming interneuron-specific expression **(suppl. Fig. 3g)**. Erbb4 RNA based on smFISH was significantly elevated in Dlx-138-sponge compared to Dlx-control sponge infected interneurons **(Fig. 3f)**, demonstrating that miR-138-5p cell-autonomously inhibits Erbb4 expression in hippocampal interneurons.

Our experiments so far suggest that impairments in interneurons are causally involved in short-term memory deficits upon miR-138 inhibition. To address the hypothesis that the observed short-term deficits are hippocampus-dependent, we stereotactically injected Dlx-138-sponge or Dlx control sponge into the dorsal hippocampus of young adult mice and assessed short-term memory 4 weeks later, as described. Specific expression of 138- and control-sponge in hippocampal interneurons was confirmed by immunostaining **(Fig. 4a)**. The previously observed 138-sponge mediated impairments in Y-maze novel preference and NOR were fully recapitulated by this approach **(Fig. 4b, 4c)**, strongly suggesting that miR-138-5p function in hippocampal interneurons is required for intact short-term memory. Importantly, short-term memory was fully preserved upon hippocampal interneuron-specific expression of the control sponge **(Fig. 4b, 4c)**, providing an independent confirmation of the specificity of our approach for miR-138-5p inactivation.

**Figure 4.**
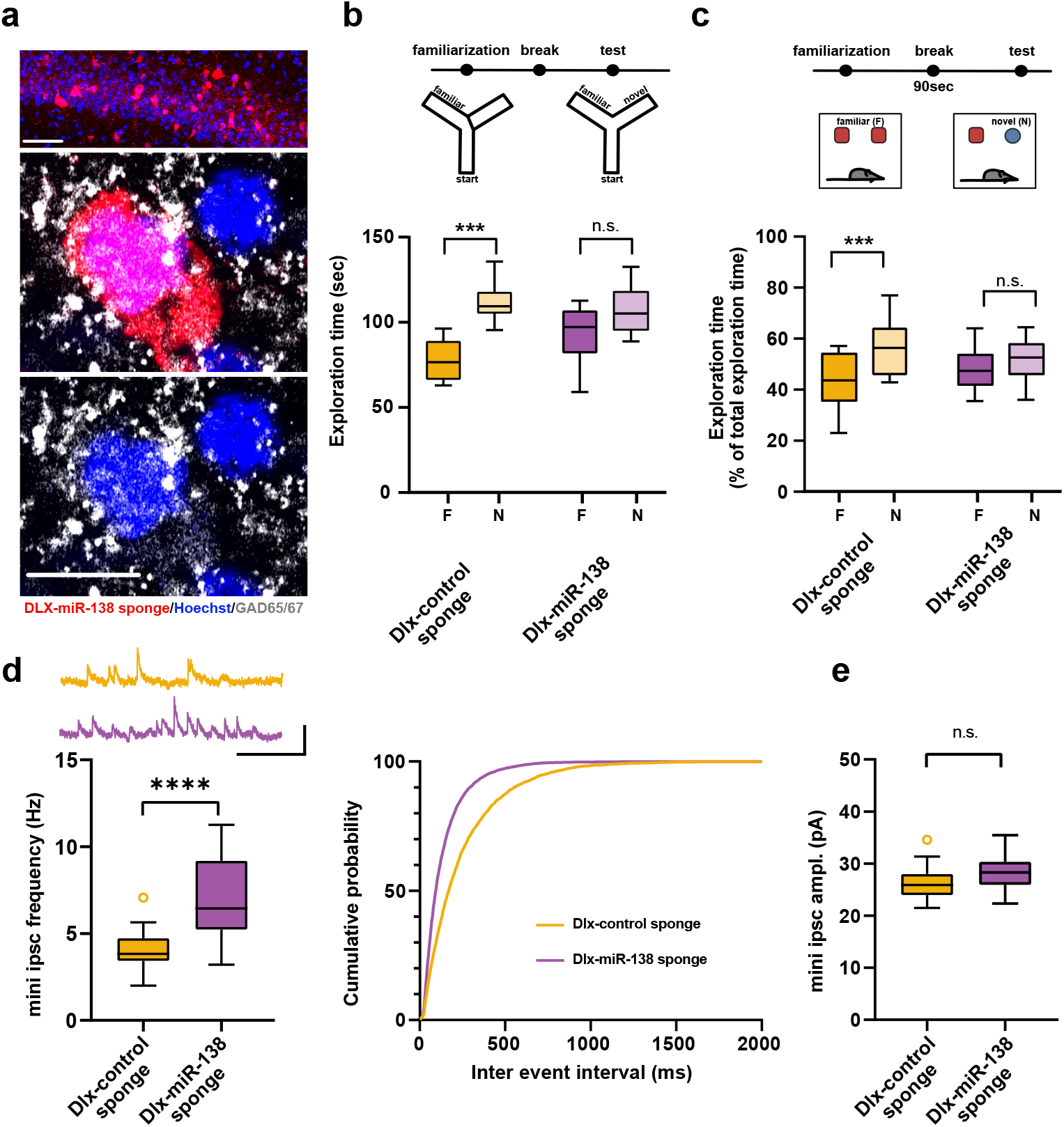
**(a)** Representative pictures of hippocampal interneurons *in vivo* infected with rAAV-Dlx-138-sponge. Upper panel: infected interneurons in hippocampal area CA1. Scale bar=50 μm. Middle and lower panel: neurons in hippocampal area CA1 at higher magnification. Left neuron: infected with rAAV-Dlx-138-sponge, expressing GAD65/67; right neuron: not infected, no GAD65/67 expression. Scale bar=10 μm. red: mCherry; grey: GAD65/67, blue: Hoechst nuclei. **(b)** Upper panel: schematic representation of the Y maze novelty preference task; lower panel: exploration time spent in familiar (F) and novel (N) arm; Dlx-control sponge n=8 mice; Dlx-miR-138 sponge n=9 mice; ****p=0.00005; n.s.=0.079 (Student’s two-tailed heteroscedastic t test). **(c)** Upper panel: schematic representation of the novel object recognition task; lower panel: exploration time presented as percentage of total time spent with either novel or familiar object; Dlx-control sponge n=9 mice; Dlx-miR-138 sponge n=10 mice; *p=0.03; n.s.=0.33 (Student’s two tailed heteroscedastic t test). **(d)** mIPSC frequency in CA1 pyramidal neurons. Upper panel left: example traces, Dlx-control sponge in orange, Dlx-miR-138 sponge in purple, scale bar: 50 pA, 200 ms. Lower panel left: mIPSC frequency (Dlx-control sponge: range, from 2.0 to 7.1 Hz; median, 3.8 Hz; IQR, 1.3 Hz. Dlx-miR-138 sponge: range, from 3.2 to 11.3 Hz; median, 6.4 Hz; IQR, 4.0 Hz; ****p < 0.0001, Student’s two-tailed heteroscedastic t test). Right panel: Cumulative distribution mIPSC frequency (p<0.0001; Kolmogorov-Smirnov test). **(e)** mIPSC amplitude in CA1 pyramidal neurons (Dlx-control sponge: range, from 21.5 to 34.6 pA; median, 25.9 pA; IQR, 4.0 pA. Dlx-miR-138 sponge: range, from 22.3 to 35.5 pA; median, 28.3 pA; IQR, 4.4 pA; n.s. p=0.13 Student’s two-tailed heteroscedastic t test). Dlx-control sponge n=19 cells/2mice; Dlx-miR-138 sponge n=19cells/2mice. **Figure 4 – source data:** this file contains the raw data on which the graphs in Fig. 4 are based.

Next, we investigated electrophysiological alterations that might underlie the observed short-term memory deficits. In hippocampal CA1 pyramidal neurons, which are the main targets of PV-positive (PV+) interneurons in the hippocampal circuit, frequency of miniature inhibitory postsynaptic current (mIPSC) was significantly increased in mice injected with the Dlx-138-sponge as compared to control sponge injected mice **(Fig. 4d)**. In contrast, amplitude **(Fig. 4e, suppl. Fig. 4a)** as well as rise- and decay time **(suppl. Fig. 4b)** of mIPSCs was unaltered.

Based on this observation, we speculated that miR-138-5p inactivation might be more robust in PV+ interneurons compared to other cell types. Thus, we again injected a miR-138 pbds sensor construct into the hippocampus of control or miR-138 sponge^Ub^ mice (cf. Fig. 1b), but now counterstained for parvalbumin (PV) to quantify miR-138 activity in PV+ and PV-cells **(suppl. Fig. 4c)**. Analysis of GFP/mCherry ratios revealed that expression of a miR-138 sponge significantly inactivated miR-138-5p in PV+ neurons but had only a marginal effect in PV-cells (which predominantly consist of excitatory pyramidal neurons). Thus, the observed lack of regulation of miR-138-5p targets in excitatory pyramidal neurons of 138-sponge^Ub^ mice is likely a result of inefficient sponge-mediated miR-138-5p inhibition in this cell type. Since miR-138-5p and alterations in PV+ interneuron function had recently been linked to schizophrenia (SCZ) (Pelkey et al., 2017; Watanabe et al., 2014), we performed a comparison between DEGs from 138-sponge^ub^ mouse hippocampus and cortical tissue of SCZ patients (Gandal et al., 2018). We found a significant overlap between genes upregulated in the 138-sponge^ub^ hippocampus and SCZ patients (p=0.00884; **suppl. Fig. 4d**). Taken together, our results suggest that miR-138-5p inactivation due to expression of a miR-138 sponge predominantly affects PV+ interneurons leading to an upregulation of synaptic genes which are deregulated in SCZ.

To elaborate on the functional role of miR-138-5p in PV+ interneurons, we generated 138-sponge^PV^ mice by crossing 138-floxed mice to PV-Cre mice (**Fig. 5a**). In 138-sponge^PV^ mice, the 6x-miR-138-sponge transcript is selectively expressed in the majority (about 90%) of PV-expressing inhibitory interneurons (PV-positive cells) (**Fig. 5b**). On a behavioral level, we found that locomotion and anxiety-related behavior, as measured in the open field and EPM tests, was unaltered in 138-sponge^PV^ mice compared to their littermate controls (**suppl. Fig. 5a-c**). However, similar as 138-sponge^ub^ mice (**Fig. 1c, d**) and mice injected with rAAV-Dlx5/6-138-sponge into the hippocampus **(Fig. 4b, c)**, 138-sponge^PV^ mice showed impairments in behavioral tasks addressing short-term memory, such as the Y-maze novelty preference (**Fig. 5c**) and NOR (**Fig. 5d**) tests. No genotype-dependent differences in spontaneous alternations in the Y-maze were observed (**Fig 5e**). Similar to 138-sponge^ub^ mice, associative long-term memory as assessed by conditional fear conditioning was not affected in 138-sponge^PV^ mice **(suppl. Fig. 5d)**. These results demonstrate that miR-138-5p activity in PV-positive interneurons is required to sustain proper short-term memory. Next, we investigated electrophysiological alterations that might underlie the observed short-term memory deficits. In hippocampal CA1 pyramidal neurons of 138-sponge^PV^ mice, similar to rAAV-Dlx5/6-138-sponge injected mice, frequency of miniature inhibitory postsynaptic current (mIPSC) was significantly increased in 138-sponge^PV^ as compared to control slices (**Fig. 6a**, **suppl. Fig. 6**), while neither amplitude, nor rise or decay time were changed (**suppl. Fig. 6b-d**). The total number of PV-positive interneurons in the hippocampus was similar between 138-sponge^PV^ and 138-floxed mice (**suppl. Fig. 6e**), suggesting that mIPSC frequency changes either result from an increased number of inhibitory presynaptic boutons synapsing onto pyramidal cells or an enhanced neurotransmitter release. To distinguish between these possibilities, we first analyzed presynaptic boutons contacting CA1 pyramidal cells by staining slices obtained from 138-floxed and 138-sponge^PV^ mice with antibodies against PV and the vesicular GABA transporter (VGAT). Our analysis revealed no significant difference in the number and intensity of PV-positive presynaptic boutons impinging onto the somata of CA1 pyramidal cells between 138-sponge^PV^ mice and their littermate controls (**Fig.6b**). To probe for changes in presynaptic release probability, we first recorded extracellularly stimulated inhibitory paired-pulse ratios (iPPRs) in CA1 pyramidal cells but did not observe differences between the two groups (**Fig. 6c**). Finally, we performed paired whole-cell recordings between presynaptic putative fast-spiking PV-positive interneurons in stratum pyramidale and postsynaptic CA1 pyramidal cells **(Fig. 6d-f; suppl. Fig. 6f)**. The analysis of unitary connections revealed increased, albeit not significant, uIPSC amplitudes (including failures of transmission) and success rates (i.e., an action potential elicits an IPSC) in 138-sponge^PV^ mice as compared to their control littermates (**Fig. 6d**). However, we did not observe significant changes in paired-pulse ratio (**Fig. 6e**). Our further analysis though revealed a significant decrease of the coefficient of variation (CV), indicative of more reliable unitary synaptic connections between PV-positive interneurons and CA1 pyramidal neurons in 138-sponge^PV^ mice (**Fig. 6f**). These findings may also explain why mIPSC frequency was found to be altered as these inhibitory currents reflect inputs of many GABAergic neurons onto a single excitatory pyramidal neuron. In conclusion, CA1 pyramidal neurons in 138-sponge^PV^ mice receive increased inhibitory GABAergic synaptic input from putative fast-spiking PV-positive interneurons without detectable changes in perisomatic inhibitory bouton density or size.

**Figure 5.**
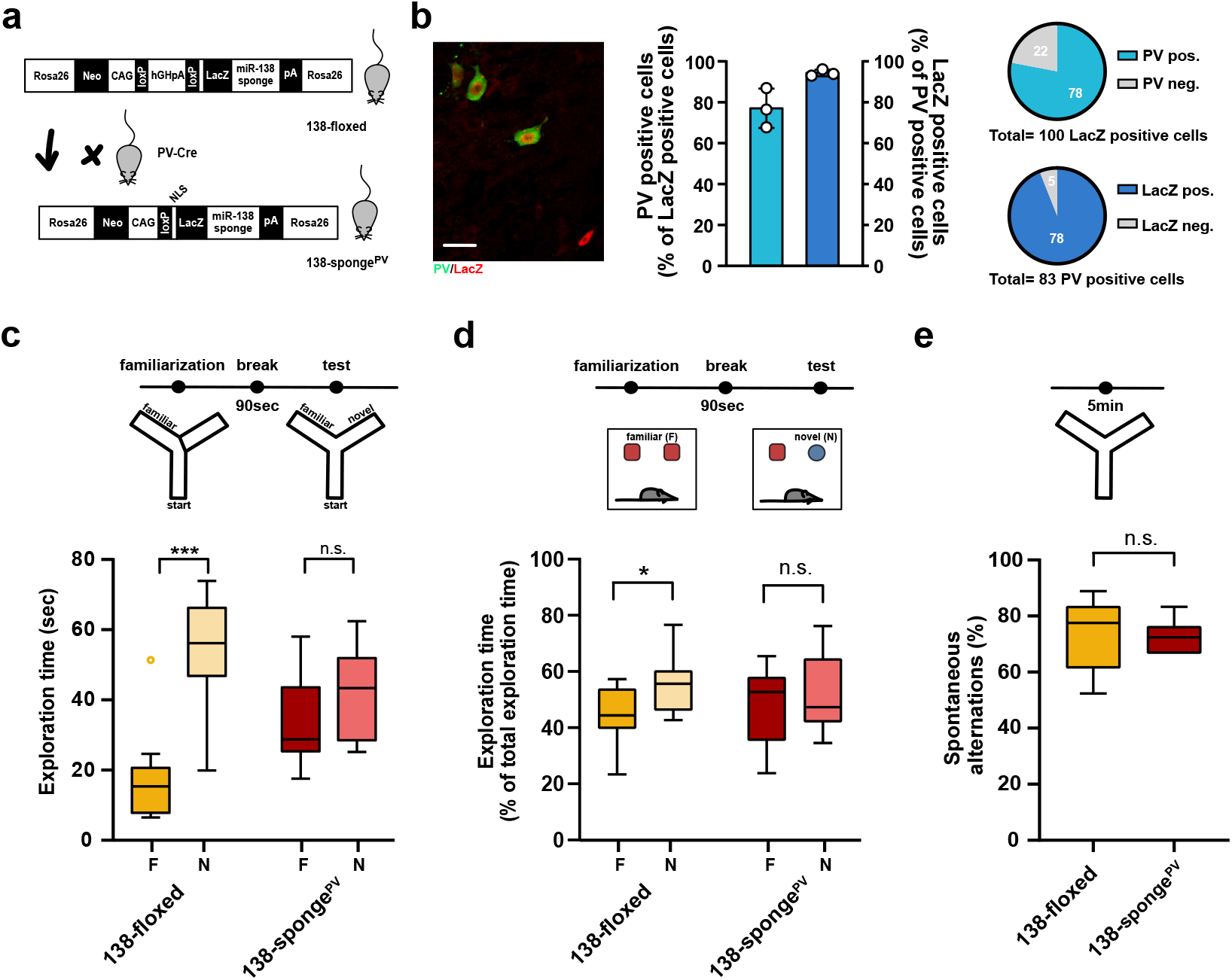
**(a)** Schematic overview of the strategy for generating miR-138 sponge^PV^ mice. **(b)** Beta-gal expression is largely restricted to PV expressing interneuron. Left panel: Representative picture from a CA1 region of a 138-sponge^PV^ hippocampal slice co-stained for the lacZ product beta-galactosidase (red) and PV (green). Scale bar=20 μm. Right panel: quantification of PV+/lacZ+ cells in 138-sponge^PV^ mice. n=3 mice. **(c)** Behavioral characterization of 138-sponge^PV^ mouse line, upper: schematic representation of the Y maze novelty preference task; lower: exploration time spent in familiar (F) and novel (N) arm; 138-floxed n=10 mice; 138-sponge^PV^ n=10 mice; ***p=0.0002; n.s. p=0.19 (Mann-Whitney test). **(d)** Upper: schematic representation of the novel object recognition task; Lower: exploration time presented as percentage of total time spent with either novel or familiar object; 138-floxed n=10 mice; miR-138 sponge n=10 mice; *p=0.035, n.s. p=0.57 (Student’s two-tailed heteroscedastic t test). **(e)** Percentage of spontaneous alternations in the Y-Maze. 138-floxed n=10; 138-sponge^PV^ n=10; n.s. p=0.90 (student’s two-tailed heteroscedastic t-test) **Figure 5 – source data:** this file contains the raw data on which the graphs in Fig. 5 are based.

**Figure 6:**
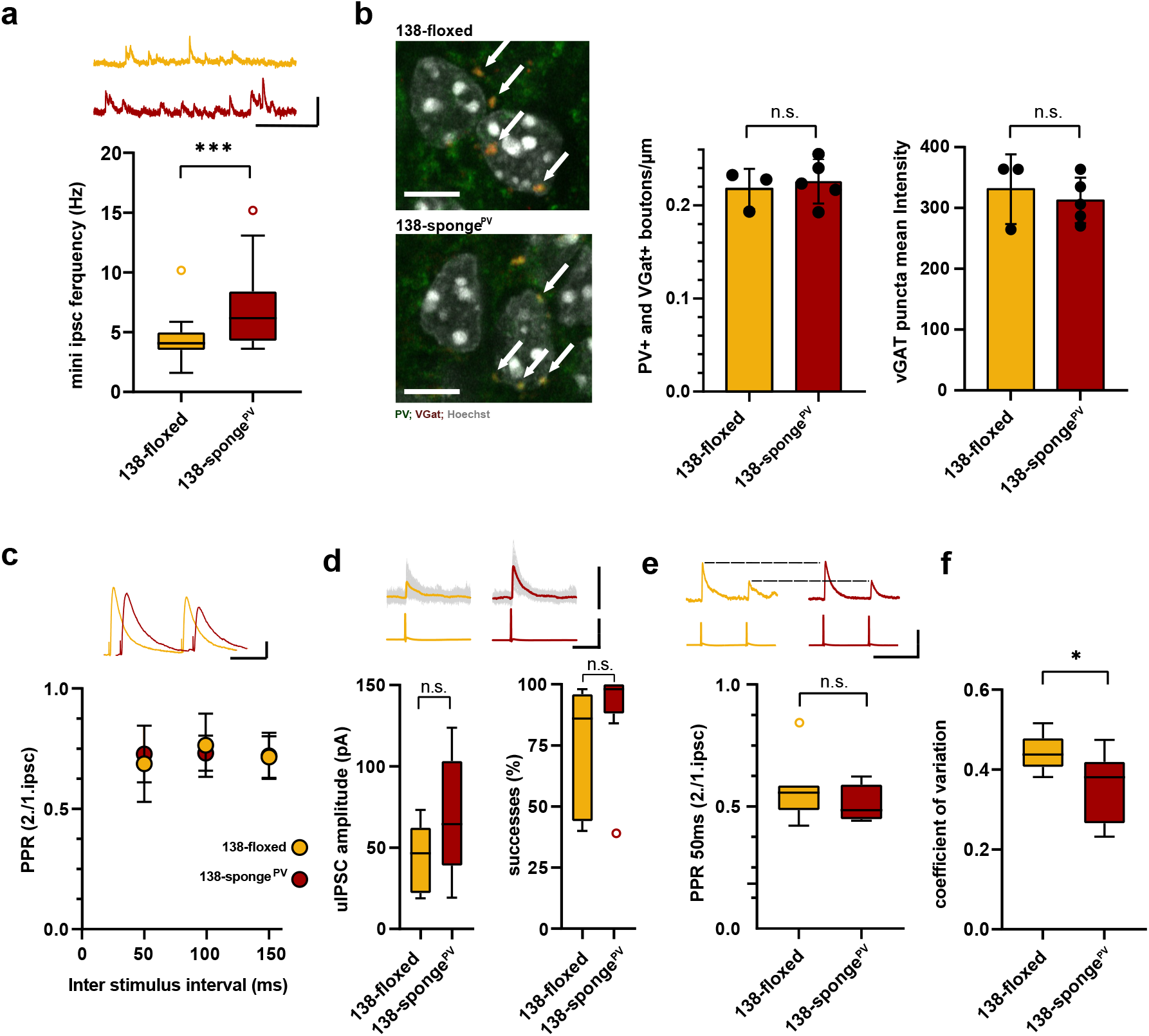
**(a)** mIPSC frequency in CA1 pyramidal neurons. Upper panel: example traces, 138-floxed in orange, 138-songe^PV^ in red, scale bar: 50 pA, 200 ms. Lower panel: mIPSC frequency (138-floxed: range, from 1.6 to 10.2 Hz; median, 4.1 Hz; IQR, 1.5 Hz. 138-sponge^PV^: range, from 3.6 to 15.2 Hz; median, 6.2 Hz; IQR, 4.2 Hz; ***p = 0.0002, Mann Whitney test). 138-floxed n=22 cells/5mice; 138-sponge^PV^ n=23cells/5mice. **(b)** PV+, VGAT+ bouton density. left panel: representative pictures (arrows point the PV+; VGAT+ boutons). middle panel: number of boutons per CA1 pyramidal neuron cell perimeter based on Hoechst counterstain. Right panel: VGAT puncta mean intensity; 138-floxed: n=73 cells/3 mice; 138-sponge^PV^: n=95 cells/5 mice; data represents the average per mouse ± s.d; n.s., p=0.65 (bouton density), n.s.=0.59 (mean intensity) (Student’s two-tailed heteroscedastic t test); **(c)** Paired pulse ratio (PPR) of stimulated IPSCs in CA1 pyramidal neurons. Upper panel: example traces of PPR (inter stimulus interval of 100ms) for 138-floxed (orange) and 138-sponge^ub^ (red); scale bar: 100 pA, 50 ms. Lower panel: PPRs for different interstimulus intervals ranging from 50 ms to 150 ms (138-floxed vs. 138-sponge^PV^ [mean ± s.d.]: 50 ms, 0.7 ± 0.2 vs. 0.7 ± 0.1 [n=13]; 100 ms: 0.8 ± 0.1 vs. 0.7 ± 0.1 [n=13]; 150 ms: 0.7 ± 0.1 [n=12] vs. 0.7 ± 0.1. n.s. p=0.45, p=0.69, and p=0.89 for 50 ms, 100 ms and 150 ms, respectively. Mann Whitney test). **(d)** Unitary connections between presynaptic fast-spiking interneurons and postsynaptic CA1 pyramidal cells. Upper panel: example traces, 138-floxed: average of 50 sweeps in orange, 26 single sweeps in grey, 138-songe^PV^: average of 50 sweeps in red, 26 single sweeps in grey; scale bar: 100 pA, 100 mV, 25 ms. Lower panel left: uIPSC amplitude (138-floxed: range, from 18.9 to 73.2 pA; median, 46.6 pA; IQR, 40.2 pA. 138-sponge^PV^: range, from 19.3 to 123.8 pA; median, 64.5 pA; IQR, 64.4 pA; n.s. p=0.11 Student’s two-tailed heteroscedastic t test). Lower panel right: Success rate (138-floxed: range, from 40 to 98 %; median, 86 %; IQR, 54 %. 138-sponge^PV^: range, from 38 to 100 %; median, 98 %; IQR, 12 %; n.s. p=0.14 Mann Whitney Test). 138-floxed n=7 pairs/3mice; 138-sponge^PV^ n=9 pairs/5mice. **(e)** Paired pulse ratio of unitary connections. Upper panel: example traces, 138-floxed in orange, 138-songe^PV^ in red, uIPSCs are normalized to the first uIPSC, scale bar: 100 mV, 50 ms. Lower panel: PPR (2^nd^/1^st^ uIPSC) (138-floxed: range, from 0.42 to 0.85; median, 0.56; IQR, 0.10. 138-sponge^PV^: range, from 0.44 to 0.62; median, 0.49; IQR, 0.14; n.s. p=0.34 Student’s two-tailed heteroscedastic t test). 138-floxed n=7 pairs/3mice; 138-sponge^PV^ n=9 pairs/5mice. **(f)** Coefficient of variation (138-floxed: range, from 0.38 to 0.52; median, 0.44; IQR, 0.07. 138-sponge^PV^: range, from 0.23 to 0.47; median, 0.38; IQR, 0.15; *p= 0.028, Student’s two-tailed heteroscedastic t test). 138-floxed n=7 pairs/3mice; 138-sponge^PV^ n=9 pairs/5mice. **Figure 6 – source data:** this file contains the raw data on which the graphs in Fig. 6 are based.

## Discussion

Here, we describe a central role for the brain-enriched miRNA miR-138-5p in the regulation of inhibitory GABAergic transmission in the hippocampus. Particularly, we find that cell type-specific inhibition of miR-138-5p in PV-positive interneurons enhances the reliability of unitary synaptic connections from putative fast-spiking PV-positive interneurons onto CA1 pyramidal neurons. The resulting increase in neurotransmitter release at this synapse is possibly due to an increase in the number of presynaptic release sites within individual boutons (Sakamoto et al., 2018), as we observed a decrease in CV, but did not detect significant changes in release probability. PV-positive interneurons are the main source of feedforward inhibition onto pyramidal neurons (Pouille and Scanziani, 2001) and have been functionally linked to working memory (Murray et al., 2011), possibly via controlling gamma oscillations (Hajos et al., 2004). We therefore propose a model whereby miR-138-5p activity in PV+ interneurons is regulating neurotransmitter release, thereby keeping pyramidal cell output in a range required for proper information processing. Since miR-138-5p controls dendritic spine morphogenesis in cultured hippocampal pyramidal neurons (Siegel et al., 2009), it might further regulate E-I balance in the hippocampal circuitry by controlling pyramidal neuron excitatory input. The absence of changes in excitatory synaptic transmission in the hippocampus of 138-sponge^ub^ mice might be due to ineffective silencing of the highly abundant miR-138-5p in pyramidal neurons via our approach. This view is supported by our observation that a miR-138-5p sensor plasmid is effectively unsilenced in PV+, but not PV-hippocampal neurons of 138-sponge^ub^ mice (suppl. Fig. 4d) and by the lack of upregulation of miR-138-5p targets preferentially expressed in pyramidal neurons (suppl. Fig. 2c). The molecular explanation for this PV-specific miR-138-5p inactivation is currently unknown.

Sponge-mediated inactivation of miRNAs is routinely used in cell culture, but studies employing this approach in transgenic mice are still scarce (Giusti et al., 2014), possibly in part because a stringent control would require the generation of an independent transgenic line. Here we addressed this issue by comparing the effects of miR-138-sponge to a highly similar, but ineffective control sponge, in primary neurons *in vitro* and in the hippocampus *in vivo* using multiple assays. Thereby, we reproducibly observed 138-sponge specific effects on reporter gene expression (suppl. Fig. 1b), dendritic spine size (suppl. Fig. 1c), miniature inhibitory postsynaptic currents (Fig. 4d, e) and short-term memory (Fig. 4b, c), strongly arguing that the effects seen in miR-138-sponge mice are due to miR-138-5p inactivation.

The molecular mechanisms downstream of miR-138-5p inactivation leading to enhanced inhibition remain to be elucidated, but likely involve upregulation of proteins organizing inhibitory synapse function. An interesting candidate in this regard represents Erbb4, which has been shown to control GABAergic transmission (Fazzari et al., 2010; Wang et al., 2018) and working memory (Tian et al., 2017; Wen et al., 2010). In particular, release probability of GABAergic synapses onto pyramidal neurons was diminished in Erbb4 KO mice (Wang et al., 2018), which is in agreement with our finding that elevated Erbb4 levels in miR-138 sponge mice are paralleled by higher mIPSC frequencies. However, additional presynaptic genes might function downstream of miR-138-5p, e.g. Rims3, which physically and functionally interacts with presynaptic voltage-dependent Ca^2+^ channels (VDCCs) and increases neurotransmitter release (Takada et al., 2015). In the future, restoring the expression of specific inhibitory presynaptic genes in miR-138 sponge mice will be needed to assess their contribution to cellular and behavioural phenotypes caused by miR-138 inhibition.

Intriguingly, interfering with miR-138-5p in inhibitory neurons of the adult hippocampus fully recapitulates deficits in inhibitory synaptic transmission and short-term memory observed in 138-sponge^Ub^ and 138-sponge^PV^ mice (Fig. 4). Although this finding does not rule out a role for miR-138-5p at early stages of development, it stresses the importance of miR-138-5p for the homeostasis of the fully developed circuitry. A similar critical role at adulthood in the regulation of GABAergic transmission and behavior was recently shown for the miR-138-5p target Erbb4 (Wang et al., 2018).

Our model of miR-138-5p being a positive regulator of memory is consistent with results obtained from previous studies performed in mice (Boscher et al., 2020; Tatro et al., 2013; Tian et al., 2018). Moreover, GWAS analysis identified miR-138-5p as a putative regulator of human memory performance (Schroder et al., 2014). A rare miR-138-2 gene variation is furthermore associated with SCZ in a Japanese population (Watanabe et al., 2014), and miR-138-5p levels are altered in the superior temporal gyrus and dorsolateral prefrontal cortex of SCZ patients (Beveridge et al., 2010) (Moreau et al., 2011). Finally, several miR-138-5p targets have been genetically linked to SCZ (e.g. *Erbb4, Drd2, Igsf9b*) (Kumar et al., 2010; Nicodemus et al., 2006) or are deregulated in the cortex of SCZ patients (e.g. *Gabra3, Fxyd6, Tmem132c*, *Baiap3*, suppl. Fig. 4d). Thus, miR-138-5p might be involved in the control of cognitive function in humans and represent a promising target for the treatment of cognitive deficits associated with schizophrenia and other neurodevelopmental disorders.

## Materials and methods

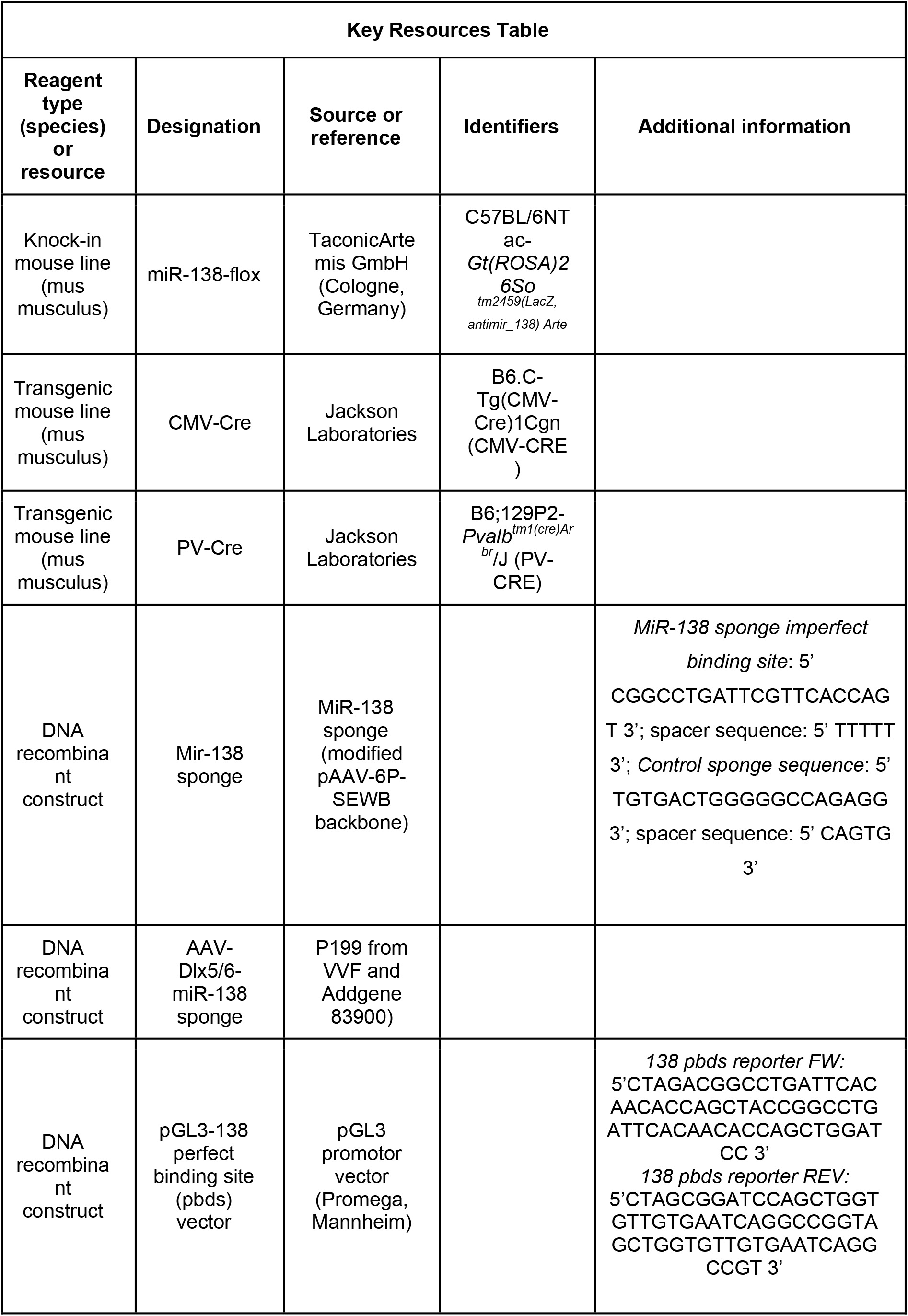
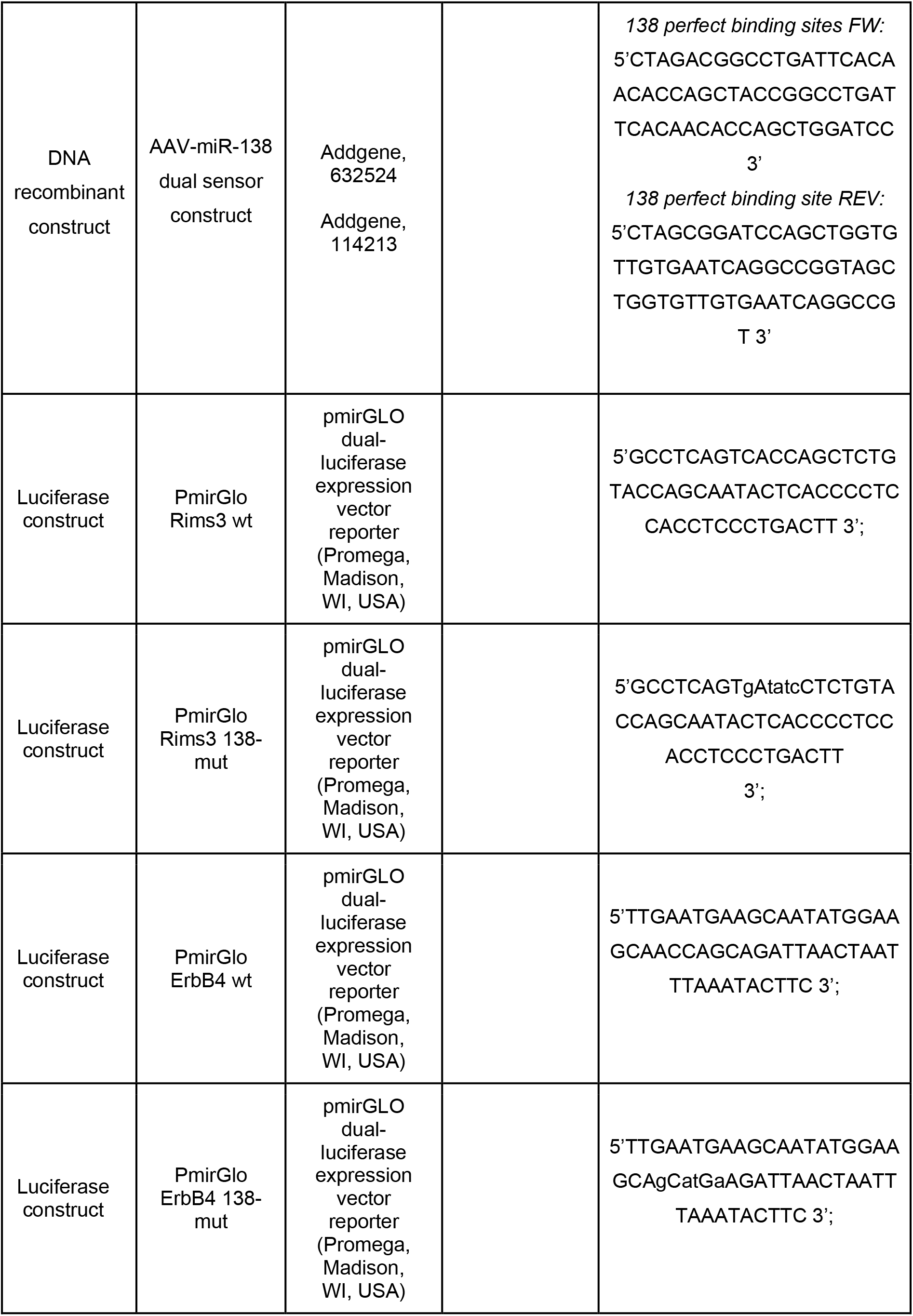
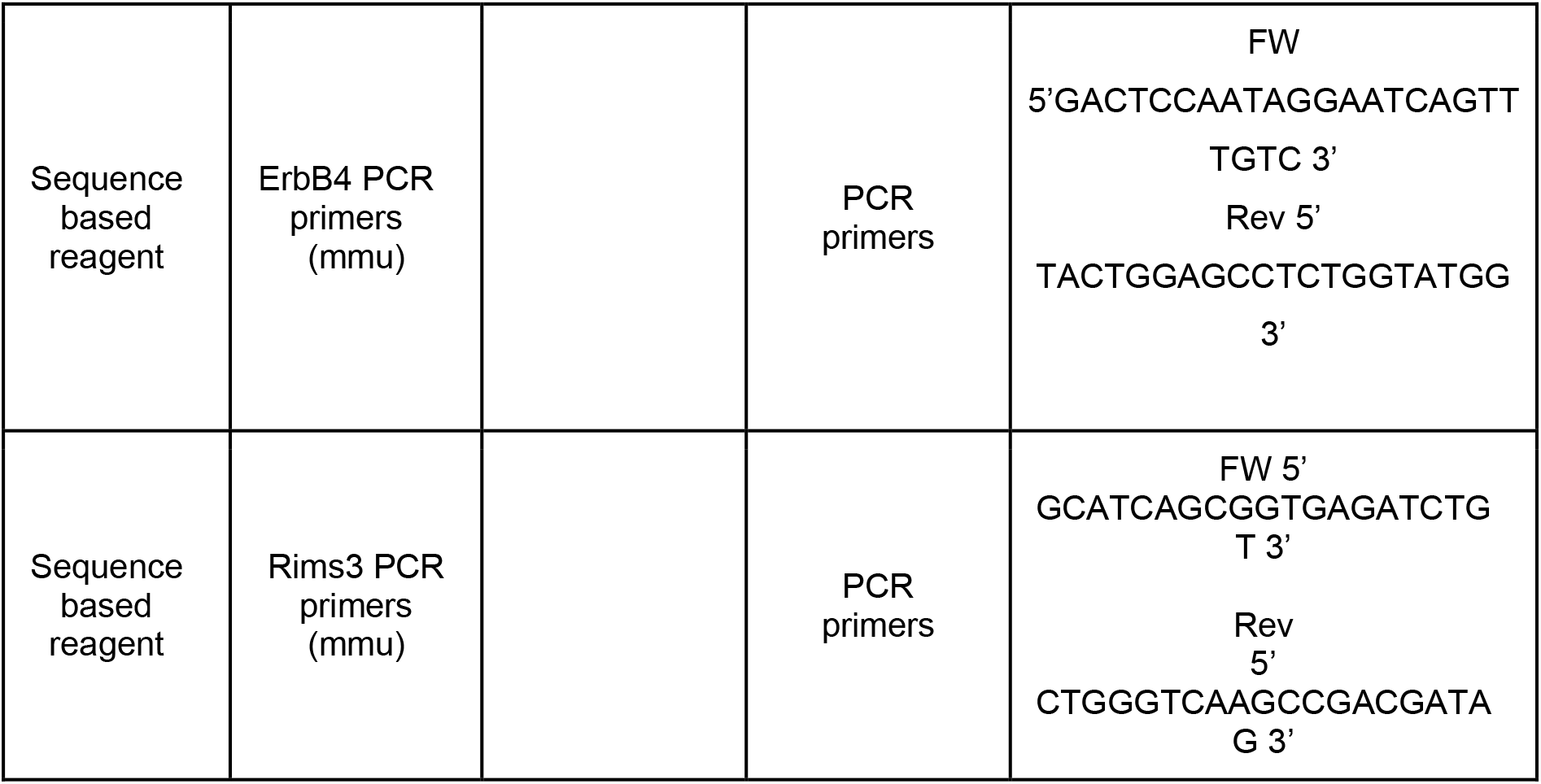

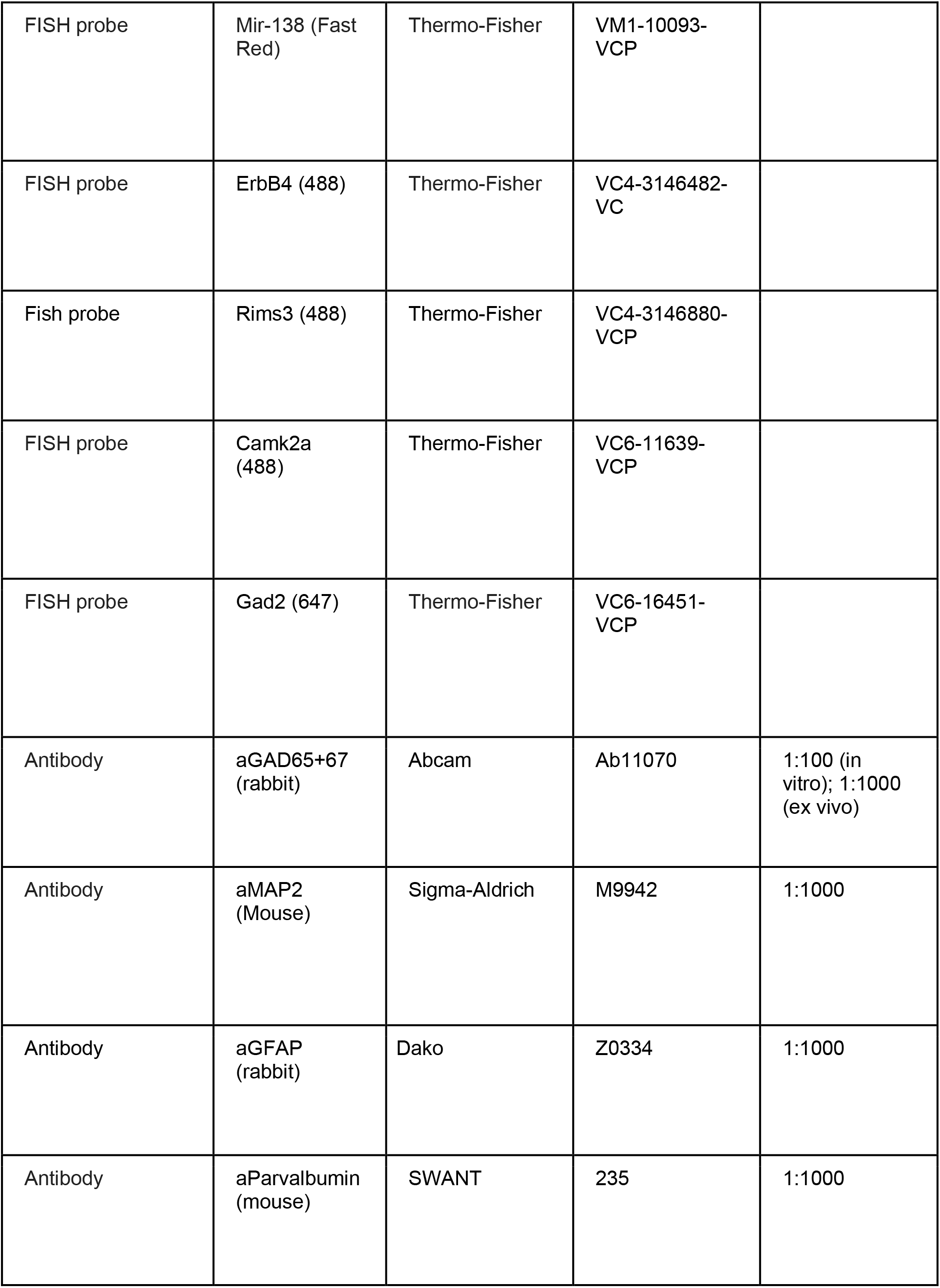
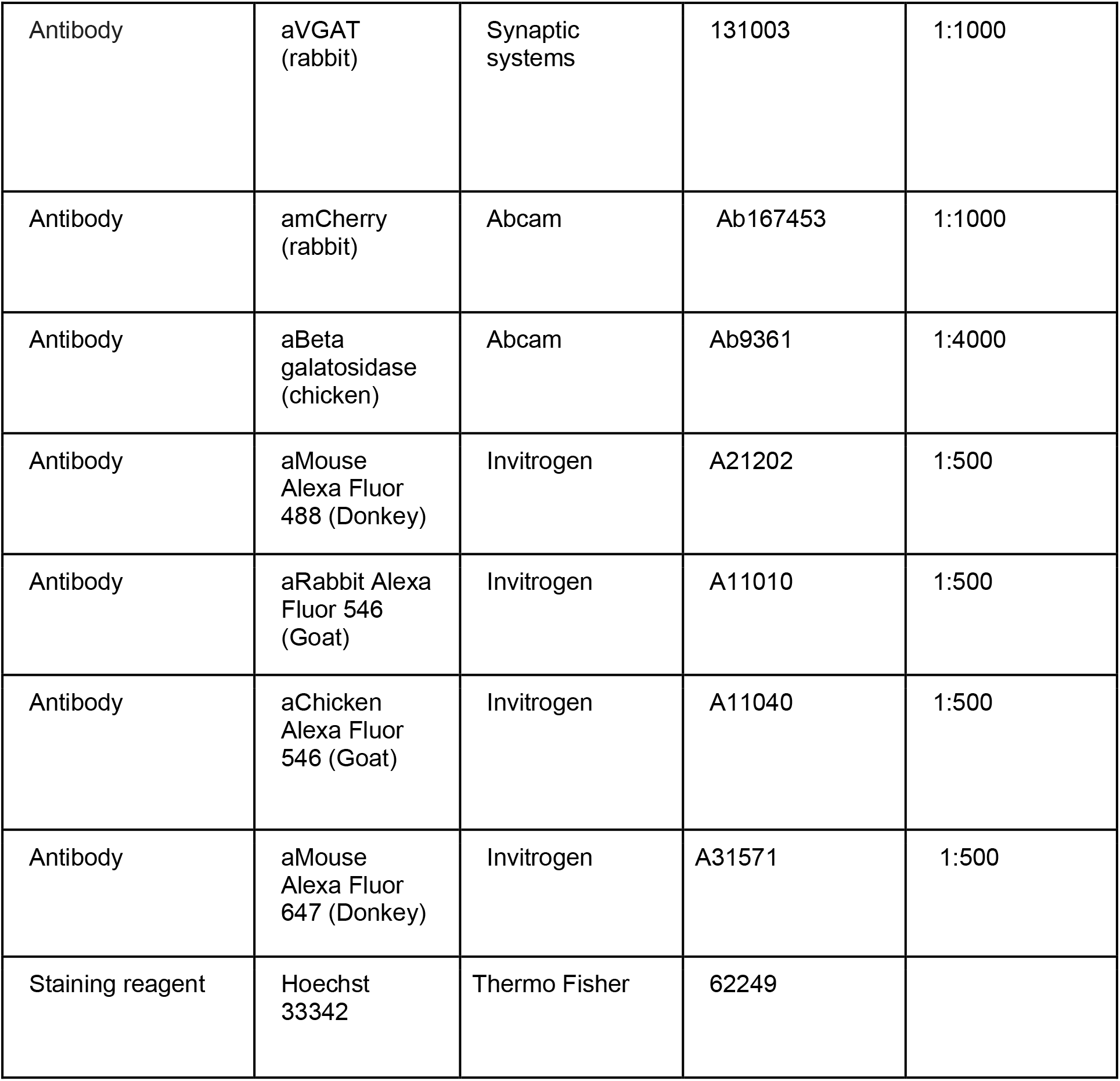

### Construct design and cloning

#### MiR-138 sponge

MiR-138 sponge was cloned into the BsrGI and HindIII restriction sites of a modified pAAV-6P-SEWB backbone, where the GFP has previously been replaced with dsRed(Christensen et al., 2010). Different number of binding sites were tested, and the plasmid containing 6 binding sites was chosen for further experiments which included the virus production for the creation of the mouse lines.

*MiR-138 sponge imperfect binding site*: 5’ CGGCCTGATTCGTTCACCAGT 3’; spacer sequence: 5’ TTTTT 3’; *Control sponge sequence*: 5’ TGTGACTGGGGGCCAGAGG 3’; spacer sequence: 5’ CAGTG 3’

#### AAV-Dlx5/6-miR-138 sponge

Dlx-138-sponge and Dlx-control-sponge were cloned into a modified mCaMKIIα-mCherry-WPRE-hGHp(A) (p199, Viral vector facility of the Neuroscience Center Zurich) backbone where 138-sponge and 138-sponge reversed (control-sponge) were previously inserted using MiR-138 sponge vector as template. The backbone was opened with MluI (NEB: R0198S) and KpnI (NEB: R0142). pAAV-mDlx-GFP-Fishell-1 (gift from Gordon Fishell; Addgene plasmid # 83900) was used to cut out Dlx enhancer and HBB promoter with MluI and EheI (NEB: R0606S).

Primers for 138 and control sponge: 5’ CACCACCTGTTCCTGTAGTGTAC 3’; 5’ CCAGAGGTTGATTATCGATAAGCTTAAC 3’.

#### AAV-CMV-Cre

AAV expressing Cre-recombinase under the control of the CMV promoter (gift from Fred Gage; Addgene plasmid #49056).

#### pGL3-138 perfect binding site (pbds) vector

The oligonucleotides were designed without any specific overhangs to allow blunt end cloning in both directions for a perfect binding site reporter and an antisense control reporter. They were annealed and ligated into a pGL3 promotor vector (Promega, Mannheim) expressing firefly luciferase. For cloning, the pGL3 vector was before digested with XbaI and the ends were blunted to insert the binding sites via blunt end cloning at the 3’end of the firefly open reading frame (ORF). After cloning, several clones were sequenced to determine sense- and antisense reporter.

*138 pbds reporter FW:* 5’CTAGACGGCCTGATTCACAACACCAGCTACCGGCCTGATTCACAACACCAGCTGGATCC 3’

*138 pbds reporter REV:* 5’CTAGCGGATCCAGCTGGTGTTGTGAATCAGGCCGGTAGCTGGTGTTGTGAATCAGGCCGT 3’

#### AAV-miR-138 dual sensor construct

An EcoR1-BglII PCR fragment spanning the pCMV promoter, mCherry coding sequence and SV40 poly adenylation signal was amplified using C1-mCherry (Addgene, 632524) as template. The PCR product was cloned into the BGlII/EcoR1 sites of the AAV-hSyn-EGFP (Addgene, 114213), upstream of the human Synapsin 1 promoter. Two miR-138 perfectly complementary binding sites were cloned into the SpeI and HIndIII restriction sites, between the EGFP coding sequence and WPRE element of AAV-hSYN-GFP, to generate the final AAV-138 sensor:

*138 perfect binding sites FW:* 5’CTAGACGGCCTGATTCACAACACCAGCTACCGGCCTGATTCACAACACCAGCTGGATCC 3’

*138 perfect binding site REV:* 5’CTAGCGGATCCAGCTGGTGTTGTGAATCAGGCCGGTAGCTGGTGTTGTGAATCAGGCCGT 3’

#### Luciferase constructs

pmirGLO dual-luciferase expression vector reporter (Promega, Madison, WI, USA) was used to clone portions of 3’ untranslated regions of the investigated mRNAs. XhoI and SalI restriction enzymes were used.

*ErbB4 wild type sequence*:

5’TTGAATGAAGCAATATGGAAGCAACCAGCAGATTAACTAATTTAAATACTTC 3’;

*ErbB4 138-mutant sequence*:

5’TTGAATGAAGCAATATGGAAGCAgCatGaAGATTAACTAATTTAAATACTTC 3’;

*Rims3 wild type sequence*: 5’GCCTCAGTCACCAGCTCTGTACCAGCAATACTCACCCCTCCACCTCCCTGACT T 3’;

*Rims3 138-mutant sequence*:

5’GCCTCAGTgAtatcCTCTGTACCAGCAATACTCACCCCTCCACCTCCCTGACTT 3’;

### Primary neuronal cell culture

Cultures of dissociated primary cortical and hippocampal neurons from embryonic day 18 (E18) Sprague-Dawley rats (Janvier, France) were prepared and cultured as described previously (Schratt et al., 2006). Animal euthanasia was approved by the local cantonal authorities (ZH196/17).

### Transfection

Transfection of primary neurons was performed using Lipofectamine 2000 (Invitrogen, Karlsruhe). For each well of a 24-well plate a total of 1 ug DNA was mixed with a 1:50 dilution of Lipofectamine in NB/NBP medium. After an incubation of 20 min at room temperature, it was further diluted 1:5 in NB/NBP medium and applied to the cells. Neurons were incubated for 2 h with the mix. A 1:1000 dilution of APV (20mM) in NB+/NBP+ was applied for 45-60 min afterwards before exchanging with NB+/NBP+.

### Luciferase assay

Luciferase assays were performed using the dual-luciferase reporter assay system on a GloMax R96 Microplate Luminometer (Promega). pmirGLO dual-luciferase expression vector reporter (Promega, Madison, WI, USA) was used to clone portions of 3’ untranslated regions of the investigated mRNAs. PcDNA3 was used to balance all amounts to a total of 1 ug DNA per condition.

### Animal lines

The C57BL/6NTac-*Gt(ROSA)26So ^tm2459(LacZ, antimir_138) Arte^* (hereafter named “138-floxed”) mouse lines was created at TaconicArtemis GmbH (Cologne, Germany). The targeting strategy allowed the generation of a constitutive LacZ-miRNA138 Sponge Knock-In (KI) allele in the C57Bl/6 mouse ROSA26 locus via targeted transgenesis. The presence of the loxP-flanked transcriptional STOP cassette is expected to terminate the transcription from the CAG promoter and thus prevent the expression of the NLS-LacZ miRNA138 Sponge cDNA, which allows this line to be used as a control (138-floxed line). The constitutive KI allele was obtained after Cre-mediated removal of the loxP-flanked transcriptional STOP cassette from the conditional KI allele, by crossing the 138-flox line with a B6.C-Tg(CMV-Cre)1Cgn (CMV-CRE) line, which allows the expression of the sponge construct (138-sponge^ub^ mice). In all experiments, heterozygous male mice were used. The 138-sponge^PV^ line was generated by crossing 138-floxed line with a previously characterized B6;129P2-*Pvalb^tm1(cre)Arbr^*/J (PV-CRE) line.

### Behavioral Experiments

Animals were housed in groups of 3-5 per cage, with food and water *ad libitum*. All experiments were performed on adult male mice (3-5 months old). The animal house had an inverted light-dark cycle, all testing was done during the dark phase (8 am-8 pm). Mice were handled for 10 minutes for 5 days before the experiments began. All measures were analyzed by Noldus Ethovision xt 14, unless stated differently. During the experimental phase, mice were transported individually and allowed to acclimatize to the experimental room in a holding cage for at least 20 minutes before the beginning of the task. At the end of each experiment they were transported back into the animal storage room in their holding cage and placed back into their original home cage with their littermates. 10ml/l detergent (For, Dr .Schnell AG) was used to clean equipment in between trials. Tasks which required the use of the same apparatus were scheduled at least 4 days apart. Two separate cohorts of mice were tested (7 mice each). No cohort-specific differences were found. The behavioral essays were performed from the least to the most stressful: home cage activity, open field, y maze, elevated plus maze, Novel object recognition, contextual fear conditioning. All animal experiments were performed in accordance with the animal protection law of Switzerland and were approved by the local cantonal authorities (ZH017/18).

#### Open field (OFT)

OFT was performed as described previously (Lackinger et al., 2019). Each session lasted 30 minutes.

#### Y maze

Spontaneous alternation: mice were placed in a Y shaped maze (8.5cm width x 50cm length x 10cm height) for 5 minutes. They were free to explore the whole maze and the alternation between the arms was calculated. Novelty preference test: mice were given 5 minutes to explore 2 arms of the maze during the familiarization phase. A door made from the same Plexiglas used for the walls was used to prevent the access of the subjects into the third arm. At the end of the familiarization phase, mice were placed in their holding cage for 90 seconds, while the apparatus was cleaned to avoid olfactory trails. Mice were then placed back into the maze for the test phase, where all three arms where accessible. Preference ratio between the familiar and new arm was then scored based on the time spent in those arms.

#### Elevated Plus Maze

The maze was elevated 60 cm from the floor, with two arms enclosed by dark Plexiglas walls (5cm width × 30cm length × 15cm height), two opposing open arms and a central platform/intersection. Experiments were conducted in a homogenously illuminated room, with the maze placed in the center of the room. Mice were placed in the central platform of the elevated plus maze. The position and motion of the animals was automatically determined and recorded for 5 min. Time spent and distance travelled in the different arms was scored.

#### Novel Object Recognition

Objects were based on the Nature protocol published previously (Leger et al., 2013). They were tested beforehand to assess that no object was preferred, and they were randomized between trials and genotypes. Subjects were placed in the open field arena with two items of the same objects for a 5 min familiarization period. After a 90 seconds break in their holding cage while the arena and the objects were cleaned and one object changed, the mice were put back in the arena for the test phase for 5 minutes. Time spent exploring the new object was scored manually. Scoring took into consideration the time spent exploring the two objects and the number of visits (nose of the experimental mouse within a 3cm radius from the object).

#### Contextual Fear Conditioning

This task was performed as previously described (Siegert et al., 2015). Briefly, mice were placed in the open field arena with Plexiglas walls and a metal grid bottom inside the multiconditioning chambers and a metal grid bottom (TSE fear conditioning system, TSE Systems, Germany). They were habituated for 3 min, then foot-shocked (2 s, 0.8 mA constant current) and returned to their home cages. After 24 h, mice were placed in the conditioning chamber. Freezing, defined as a total lack of movement except for heartbeat and respiration, was scored for a 3 min period.

### RNAseq

#### RNA extraction

Adult male mice (4 months old) were anaesthetized with isoflurane (Baxter, Unterschleißheim, Germany) and then quickly cervically dislocated and decapitated. Hippocampal tissue was dissected and freshly processed (RNA was extracted using mirVana microRNA Isolation Kit (Life Technologies) according to the manufacturer’s instructions.

Stranded polyA+ enriched RNA sequencing libraries were prepared at the GENCORE (EMBL, Genomics Core Facility, Heidelberg, Germany) and sequenced on an Illumina HiSeq 2000 machine using a 50nt paired-end protocol (Lackinger et al., 2019).

We removed sequencing adapter and quality-trimmed all short reads from the 3’end using FLEXBAR 2.5.3 and a Phred score cutoff >10. All reads longer than 18bp were retained and rRNA reads were subtracted in silico using bowtie2 and an index of the complete repeating rRNA unit (BK000964.3). We used the STAR aligner 2.4.2a4 to map against the mouse genome (EnsEMBL 79 genome + annotations). We performed differential gene expression analysis using Cuffdiff 2.2.1.

### Quantitative real-time PCR

qPCR was performed as described earlier (Valluy et al., 2015). Primer sequences are given below.

ErbB4 FW: 5’GACTCCAATAGGAATCAGTTTGTC 3’

ErbB4 REV: 5’ TACTGGAGCCTCTGGTATGG 3’

Rims3 FW: 5’ GCATCAGCGGTGAGATCTGT 3’

Rims3 REV: 5’ CTGGGTCAAGCCGACGATAG 3’

### Imaging

Image acquisition was done with the experimenter being blinded to the different conditions. Pictures were taken on a confocal laser scanning microscope equipped with an Airyscan detector (LSM880, Zeiss). Image analysis was carried out on Fiji (ImageJ).

*<u>PV bouton count</u>*: a 63x oil objective was used to take images from CA1 hippocampal region immunostained with PV, VGAT and Hoechst was used to counterstain the nuclei (further details in the immunostaining section). The number of PV+ and VGAT+ en passant boutons around the nuclear perimeter were counted and normalized to the total length of the perimeter.

vGAT puncta mean intensity:

vGAT puncta mean intensity: ImageJ was used to measure the intensity of all the vGAT puncta in the pictures used to quantify PV boutons.

*<u>PV cell quantification</u>*: a 20x objective was used and PV+ cells were counted on a maximum projection intensity of the CA1 region immunostained with PV (Hoechst was used to counterstain nuclei; further details in the immunostaining section). The number of PV+ cells was normalized to a defined region of interest (230μm x 460μm).

*<u>Viral mir-138 sensor</u>*: tilescans were taken with a 20x objective of the infected hippocampal region. Intensity of GFP signal was normalized on the mCherry signal per cell. For the analysis, either all neurons (Fig. 1b) or only PV+ neurons (suppl. Fig. 4d) within the CA1 region were taken into consideration.

### Spine analysis

*<u>In vitro</u>*: Hippocampal neurons were transfected at DIV 10 with 200 ng of GFP and the indicated amount of either miR-138-6x-sponge, CTR sponge, AAV backbone or the Cholesterol-modified 2’O-Me-oligonucleotides (’antagomirs’, Thermo Scientific, Karlsruhe. To balance all amounts to a total of 1 ug DNA per condition, PcDNA3 was used. At 18 DIV Cells were fixed using 4 % PFA for 15 min and mounted on glass slides using Aqua-Poly/Mount (Polysciences Inc., Eppelheim). The experimenter was blinded to all the conditions. The z-stack images of GFP-positive neurons exhibiting pyramidal morphology were taken with the 63x objective of a confocal laser scanning microscope (Carl Zeiss, Jena). Eight consecutive optical sections of the dendrites were taken at a 0.4 μm interval with a 1024 x 1024 pixel resolution. Spine volumes were analyzed with the ImageJ software using the maximum intensity projections of the z-stack images. The GFP intensity of 150-200 spines per cell was measured and normalized to the total intensity of the dendritic tree. For each experimental condition, at least 18 representative neurons derived from three independent experiments were analyzed.

*<u>In vivo</u>:* brains from 3 months old 138-floxed and 138-sponge^ub^ mice were processed with FD Rapid GolgiStain Kit (Gentaur GmbH, PK401) according to the manufacturer’s protocol. Pictures were taken with a Zeiss axio-observer 7 inverted wide field fluorescence microscope equipped with a Axiocam 702 mono Zeiss camera with an 100x oil objective. Dendritic spines were manually analyzed with Fiji (ImageJ).

### Electrophysiology

Hippocampal slices (300 μm thick) were prepared at 4 °C, as previously described (Winterer et al., 2019), from 138-floxed, 138-sponge^ub^ and 138-sponge^PV^ mice (age: 6-8 weeks) and incubated at 34°C in sucrose-containing artificial cerebrospinal fluid (sucrose-ACSF, in mM: 85 NaCl, 75 sucrose, 2.5 KCl, 25 glucose, 1.25 NaH2PO4, 4 MgCl2, 0.5 CaCl2, and 24 NaHCO3) for 0.5 h, and then held at room temperature until recording.

Whole cell patch clamp recordings were performed at 32 °C on an upright microscope (Olympus BX51WI) under visual control using infrared differential interference contrast optics. Data were collected with an Axon MultiClamp 700B amplifier and an Digidata 1550B digitizer and analyzed with pClamp 11 software (all from Molecular Devices). Signals were filtered at 2 kHz for miniature EPSCs and miniature IPSCs and digitized at 5 kHz. Stimulus evoked and unitary postsynaptic currents were filtered at 4 kHz, fast-spiking interneurons at 10 kHz, both were digitized at 100 kHz. Recording pipettes were pulled from borosilicate capillary glass (Harvard Apparatus; GC150F-10) with a DMZ-Universal-Electrode-Puller (Zeitz) and had resistances between 2 and 3 MΩ.

The extracellular solution (ACSF) was composed of (in mM) 126 NaCl, 2.5 KCl, 26 NaHCO_3_, 1.25 NaH_2_PO_4_, 2 CaCl_2_, 2 MgCl_2_ and 10 glucose. For excitatory postsynaptic currents (EPSCs) measured in CA1 pyramidal cells and for fast-spiking interneurons the intracellular solution was composed of 125 K-Gluconate, 20 KCl 0.5 EGTA, 10 HEPES, 4 Mg-ATP, 0.3 GTP and 10 Na_2_-phosphocreatine (adjusted to pH 7.3 with KOH). Cells were held at -70mV for miniature EPSC (mEPSCs) and at - 60mV for stimulus evoked excitatory postsynaptic currents (eEPSCs). 1μM Gabazine was added to isolate AMPA-mediated postsynaptic currents, additionally 1μM TTX for mEPSCs. For inhibitory postsynaptic currents (IPSCs) intracellular solution was composed of 135 Cs-Gluconate, 5 KCl, 2 NaCl, 0.2 EGTA, 10 HEPES, 4 Mg-ATP, 0.3 GTP and 10 Na_2_-phosphocreatine (adjusted to pH 7.3 with CsOH). Cells were held at +10mV for miniature IPSC (mIPSCs) and at -10mV for stimulus evoked and unitary inhibitory postsynaptic currents (eIPSCs and uIPSCs). 25μM AP-5, 10μM NBQX and 10μM SCH 50911 were added to isolate GABAa-mediated postsynaptic currents for eIPSCs, additionally 1μM TTX for mIPSCs. Synaptic currents were evoked by mono-polar stimulation with a patch pipette filled with ACSF and positioned in the middle of CA1 stratum radiatum for eEPSCs and in stratum pyramidale for eIPSCs. Series resistance of CA1 pyramidal neurons (not compensated; range, from 7.0 to 19.7 MΩ; median, 11.3 MΩ; IQR, 2.4 MΩ) was monitored and recordings were discarded if series resistance changed by more than 20%. Membrane potentials were not corrected for liquid junction potential.

For paired recordings, whole cell configuration was first established in putative fast-spiking interneurons. Cells were selected based on morphological appearance in stratum pyramidale of hippocampal CA1, on fast-spiking properties and on input resistance characteristic for PV-positive interneurons (Que et al., 2021)(**suppl. Fig. 6f**). Subsequently whole cell recordings were made from postsynaptic CA1 pyramidal neurons residing in close proximity to the presynaptic fast-spiking interneuron. Presynaptic fast-spiking interneurons were held in current-clamp mode and series resistance was compensated with the automatic bridge balance of the amplifier. The mean resting potential of the presynaptic fast-spiking interneurons was -65 ± 4.1mV (mean ± sd; **suppl. Fig. 6f)**. Fast-spiking interneurons were stimulated at 1Hz for uIPSCs and at 0.03Hz for paired pulse uIPSCs. To characterize the discharge behavior of fast-spiking interneurons, depolarizing steps (50pA) of 1500ms were applied. The spiking frequency (**suppl. Fig. 6f**) was determined at 800 to 1000pA current injection. Input resistance was calculated from the mean deviation from baseline of steady state voltage responses evoked by -150, -100 and -50 pA current injections.

### Cell type enrichment analysis

For the cell type enrichment analysis, we used the single-cell data and annotation from Zeisel et al. (2018). We downloaded single-cell count data and annotation from https://storage.googleapis.com/linnarsson-lab-loom/l5_all.loom, restricted it to hippocampus cells, and aggregated to the pseudo-bulk level using muscat (Crowell et al., 2020) and the authors’ cell identities. We retained only cell types represented by more than 50 cells, and normalized using TMM (Robinson and Oshlack, 2010). For each gene, we then identified the cell type in which it was the most highly expressed, forming gene sets for each cell type. We then created a sponge differential expression signature by multiplying the sign of the foldchange with the - log10(p-value) of each gene and looked for enrichment of gene sets in this signature using fgsea (Korotkevich et al., Biorxiv 2021, doi: 10.1101/060012).

### GO enrichment analysis

The R Bioconductor package TopGo (v.2.42.0) was used to perform Gene Ontology enrichment analysis, essentially as in (Lackinger et al., 2019).

To sum up, genes were annotated with the “Cellular Component” ontology and significantly changed genes subsequently tested against the expressed background. For the main figure we plotted the Top10 GO-Terms ranked by significance (filtering out those with more than 200 annotated genes).

### miRNA binding site analysis

Mouse 3’UTR positions of predicted conserved microRNA binding sites were downloaded from Targetscan (version 7.2)(Agarwal et al., 2015) and filtered for the seed of miR-138-5p. Putative miR-138 targets (including site-type information) were then aligned with the genes and log fold changes (logFC) as obtained by the differential expression analysis. Plotted is the cumulative proportion (logFC rank (-1) / number of genes (-1)) over the logFC.

### Single-molecule fluorescence *in situ* hybridization (smFISH)

smFISH for miRNA detection on hippocampal neuron cultures was performed using the QuantiGene ViewRNA miRNA Cell Assay kit (Thermo Fisher) according to the manufacturer’s protocol with slight modifications. To preserve dendrite morphology, protease treatment was reduced to a dilution of 1:10,000 in PBS for 45 sec. smFISH for mRNA detection was performed using the QuantiGene ViewRNA ISH Cell Assay kit (Thermo Fisher) as previously described (Valluy et al., 2015), but omitting the protease treatment. After completion of the FISH protocol, cells were washed with PBS, pre-blocked in gelatin detergent buffer and processed for immunostaining. Pictures represent maximum intensity projections of z-stack images taken on a confocal laser scanning microscope equipped with an Airyscan detector (LSM880, Zeiss).

### Stereotactic surgery

The viral vector (aaAAV-9/2 [hCMV-mCherry-SV40p(A)]rev-hSyn1-EGFP-2x (miR-138-5p)-WPRE-SV40p(A)) was produced by the local Viral Vector Facility (VVF9 of the Neuroscience Center Zurich). The produced virus had a physical titer of 6.1 x10e^12^ vector genomes/ml. Stereotactic brain injections were performed on two- to three-month-old 138-floxed and 138-sponge^ub^ mice as previously described(Zerbi et al., 2019)). Briefly, mice were anesthetized with isoflurane and subsequentially placed onto the stereotaxic frame. Before and after the procedure, animals received subcutaneous injection of 2 mg/kg Meloxicam for analgesia and local anaesthetic (Emla cream 5% lidocaine, 5% prilocaine) was distributed on the head. Animals were injected bilaterally with 1µl of virus into the dorsal hippocampus (coordinates from bregma: anterior/posterior −2.1 mm, medial/lateral ± 1.5 mm, dorsal/ventral −1.8 mm) and 1µl of virus in the ventral hippocampus (coordinates from bregma: anterior/posterior −3.3 mm, medial/lateral ± 2.75 mm, dorsal/ventral −4.0 mm). Post-operative health checks were carried on over the three days after surgery.

### Tissue collection

Animals were sacrificed by intraperitoneal injection of pentobarbital (150 mg/kg). When in deep anaesthesia, mice were perfused intracardially with ice-cold PBS pH 7.4, followed by perfusion with 4% paraformaldehyde in PBS pH 7.4. The brains were then isolated and postfixed for 2-3h (138-floxed; 138-sponge^PV^ mice) or overnight (138-flox and 138-sponge^ub^ mice) at 4°C. The fixed tissue was placed in sucrose solution (30% sucrose in PBS) for 24h and frozen in tissue mounting medium (OCT mounting media, VWR chemicals). The tissue was coronally sectioned at 50-60µm thickness on a cryostat, immediately placed in ice-cold PBS and subsequentially conserved in cryoprotectant solution (15% glucose, 30% ethylene glycol, 5mM NaH_2_PO_4_*H_2_O, 20mM Na_2_HPO_4_*2H_2_O) at -20°C.

### Immunohistochemistry

Ex *vivo (cryosections)*: For immunofluorescence, cryosections were washed in ice-cold PBS for 30 minutes, placed on microscope slides (Menzel-Gläser SUPERFROST PLUS, Thermo Scientific) and air-dried for 5-10 min. Afterwards, permeabilization was performed by incubating sections in permeabilization solution (0.5% Triton X-100 in PBS) for 30 min at room temperature, followed by a blocking step in blocking buffer (0.5% Triton X-100, 300mM NaCl, 10% Normal Goat Serum in PBS) with addition of blocking Reagent (VC-MKB-2213, Adipogen Life Sciences, 1:10 dilution) for 1h at room temperature. Cryosections were washed in PBS twice for 10 min at room temperature and incubated with primary antibodies in blocking buffer overnight at 4°C. Subsequentially, the sections were washed 3 times with blocking buffer, incubated with secondary antibodies and Hoechst 33342 in blocking buffer for 1h at room temperature, washed again three times in blocking buffer, washed in PBS, air-dried and mounted with Aqua-Poly/Mount (POL18606, Polysciences).

*Ex vivo (vibratome sections)*: For immunofluorescence, 300µM sections were sliced on a vibratome in ice cold sucrose-containing artificial cerebrospinal fluid, and then fixed in 4% PFA overnight. Free-floating sections were washed in ice-cold 1X PBS for 30min, permeabilized by incubation in 0.5% PBX (0.5% Triton X-100 in PBS) for 30 min at room temperature, followed by a quenching step in 0.1M Glycine to reduce background autofluorescence. Sections were blocked in blocking buffer (0.5% PBX, 10% normal goat serum in PBS) for 1h at room temperature, and then incubated in primary antibodies for 36 hours at 4°C. Sections were washed in 0.05% PBX twice for 10 min at room temperature, and then incubated with secondary antibodies in 0.05% PBX with 10% normal goat serum at room temperature for 3 hours.

Subsequentially, the sections were washed 3 times with 0.05% PBX, incubated with Hoechst 33342 in 1X PBS for 10min, and then washed again three times in PBS, mounted on SuperFrost+ slides with Aqua-Poly/Mount (POL18606, Polysciences). A 40uM Z-stack (with 0.25uM z-step) was acquired on a Zeiss LSM 880 confocal microscope, with a Plan-Apochromatic 63X/1.4NA DIC M27 oil objective and FastAiryScan detector settings in the hippocampal CA1 region.

In *vitro*: cells were washed once with NBP, fixed for 20 minutes with 4%PFA and, after five washes with PBS (3xfast;2×5minutes), permeabilized with GDB (20 mM sodium phosphate buffer (pH 7.4), 450 mM NaCl, 0.3 % Triton X-100, 0.1 % gelatin, ddH_2_O) for 20 minutes. Primary antibody was diluted in GDB and incubated for 1h at room temperature, followed by five washes with PBS (3×fast;2×5minutes). Secondary antibody was diluted in GDB and incubated in the dark, at room temperature for 1h. The cells were then washed again with PBS (3×fast;2×5minutes). In the second last wash Hoechst 33342 was added to the PBS. The coverslips were mounted with Aqua-Poly/Mount (POL18606, Polysciences) on microscope slides (Menzel-Gläser SUPERFROST PLUS, Thermo Scientific).

### Statistics

Statistical analysis was performed on either GraphPad Prism 8.0 or RStudio. Plots were generated in R, mainly using the packages ggplot2, ggsci, ggrepel and scales. For data sets with n>4, Box plots (Tukey style) were used. For data sets with n<5, average +/- s.d., including individual data points, is shown. Normal distribution of data was tested with the Shapiro-Wilk test. Based on the results, parametric (e.g. Student’s t-test) or non-parametric (e.g. Mann-Whitney, Kolmogorov-Smirnov) tests were used.

The detailed parameters (n, p-value, test) for the statistical assessment of the data are provided in the figure legends.

## Acknowledgments

We are grateful to S. Brown and M. Müller for valuable comments on the manuscript, T. Wüst and C. Furler for excellent technical assistance, T. Germade for help with bioinformatics, D. Colameo for help with animal perfusions and microscopy data analysis scripts, A. Loye for help with histology and T. Demeter for help with cloning. We thank G. Fishell, F. Gage and the Viral Vector Facility of the Neuroscience Center Zurich for sharing plasmids. This work was supported by grants from the DFG (SCHR 1136/4-2) and the ETH (24 18-2 (NeuroSno).

## Data and code availability

RNA-seq data (Fig. 2, suppl. Fig.2) has been deposited to GEO (accession no. GSE173982). Source data are attached as zip folders to the individual figures of the submission.

---

**Supplementary Figure 1.**
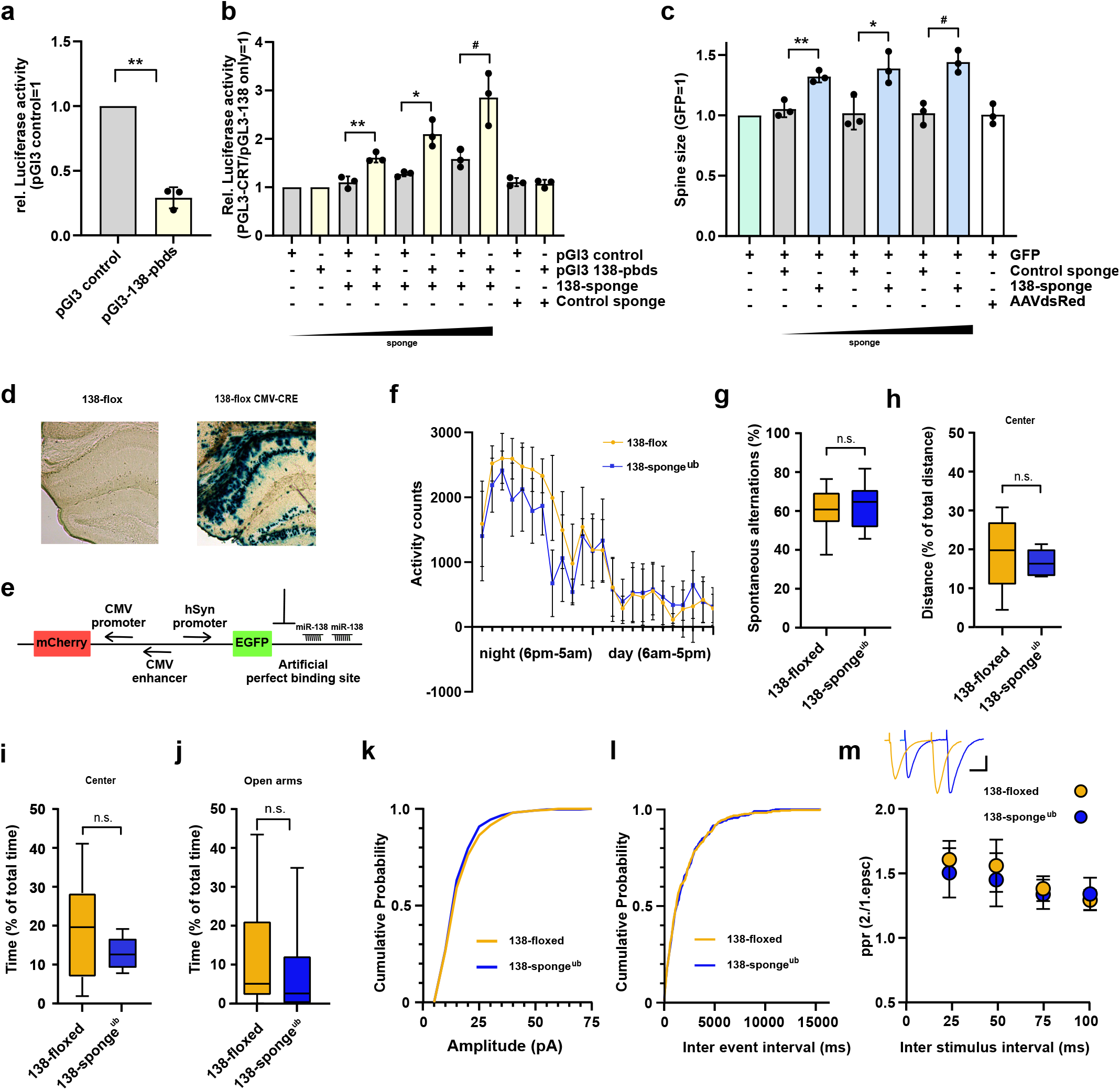
**(a)** Relative luciferase activity in hippocampal neurons (DIV 12-17) transfected with pGL3 CTR (control) or pGL3-138 pbds (sensor) constructs. pGL3-CTR=1. n=3 independent transfections. **p=0.0013 (One sample t test). **(b)** Relative luciferase activity in hippocampal neurons (DIV 12-17) transfected as in a), in addition with either control (100ng) or increasing amounts of 138 sponge (25-100ng). pGL3-CTR/pGL3-138 only = 1. n=3 independent transfections, **p=0.005, *p=0.032, #p=0.045 (student’s two-tailed heteroscedastic t test). **(c)** Quantification of relative dendritic spine volume in rat hippocampal neurons (DIV10-18) transfected with GFP and increasing amounts (25-100ng) of either control or 138 sponge. GFP only = 1. N = 3 independent transfections; each value represents at least 150 spines per cell, from 6 individual neurons per experiment. **p=0.006, *p=0.027, #p=0.004 (student’s two-tailed heteroscedastic t-test). **(d)** Representative enzymatic b-Gal staining of hippocampal slices obtained from 138-floxed mice (P21) injected with either saline (left) or rAAV-CMV-CRE (right). **(e)** Schematic of the sensor principle. **(f)** Activity counts of mice from indicated genotypes monitored over 24 h in their home cage; 138-floxed: n=7 mice, 138-sponge^ub:^ n=7 mice; data shown as mean ± s.d. **(g)** Percentage of spontaneous alternations in the Y-Maze. 138-floxed n=14 mice; 138-sponge^ub^ n=14 mice; p=0.43 (student’s two-tailed heteroscedastic t-test). **(h, i)** Percentage of total distance travelled(h)/time spent(i) in the center of an open field arena during 30 min exploration by mice of the indicated genotypes; 138-floxed n=14 mice; 138-sponge^ub^ n=14 mice; (h) p=0.30, (i) p=0.054 (Student’s two-tailed heteroscedastic t test). **(j)** Percentage of total time spent in the open arms of an elevated plus maze (EPM) during 5 min exploration by mice of the indicated genotypes; 138-floxed: n=14 mice; 138-sponge^ub^:n=14 mice; p=0.50 (Mann-Whitney test). **(k, l)** Cumulative distribution mEPSC amplitude (p=0.65; Kolmogorov-Smirnov test) (k) and frequency (p=0.85; Kolmogorov-Smirnov test) (l). **(m)** PPR of stimulated EPSCs in CA1 pyramidal neurons. Upper panel: example traces of PPR (inter stimulus interval of 50ms) for 138-floxed (orange) and 138-sponge^ub^ (blue); scale bar: 100 pA, 20 ms. Lower panel: PPRs for different interstimulus intervals ranging from 25 ms to 100 ms (138-floxed vs. 138-sponge^ub^ [mean ± s.d.]: 25 ms, 1.6 ± 0.1 vs. 1.5 ± 0.2 [n=13]; 50 ms: 1.6 ± 0.2 vs. 1.5 ± 0.2 [n=13]; 75 ms: 1.4 ± 0.1 vs. 1.3 ± 0.1 [n=11]; 100 ms: 1.3 ± 0.1 vs. 1.3 ± 0.1 [n=11]. n.s. p=0.14, 0.09, 0.17 and 0.13 for 25, 50, 75 and 100 ms inter stimulus intervals, respectively. Mann-Whitney test.) **Suppl. Figure 1 – source data:** this file contains the raw data on which the graphs in suppl. Fig. 1 are based.

**Supplementary Figure 2.**
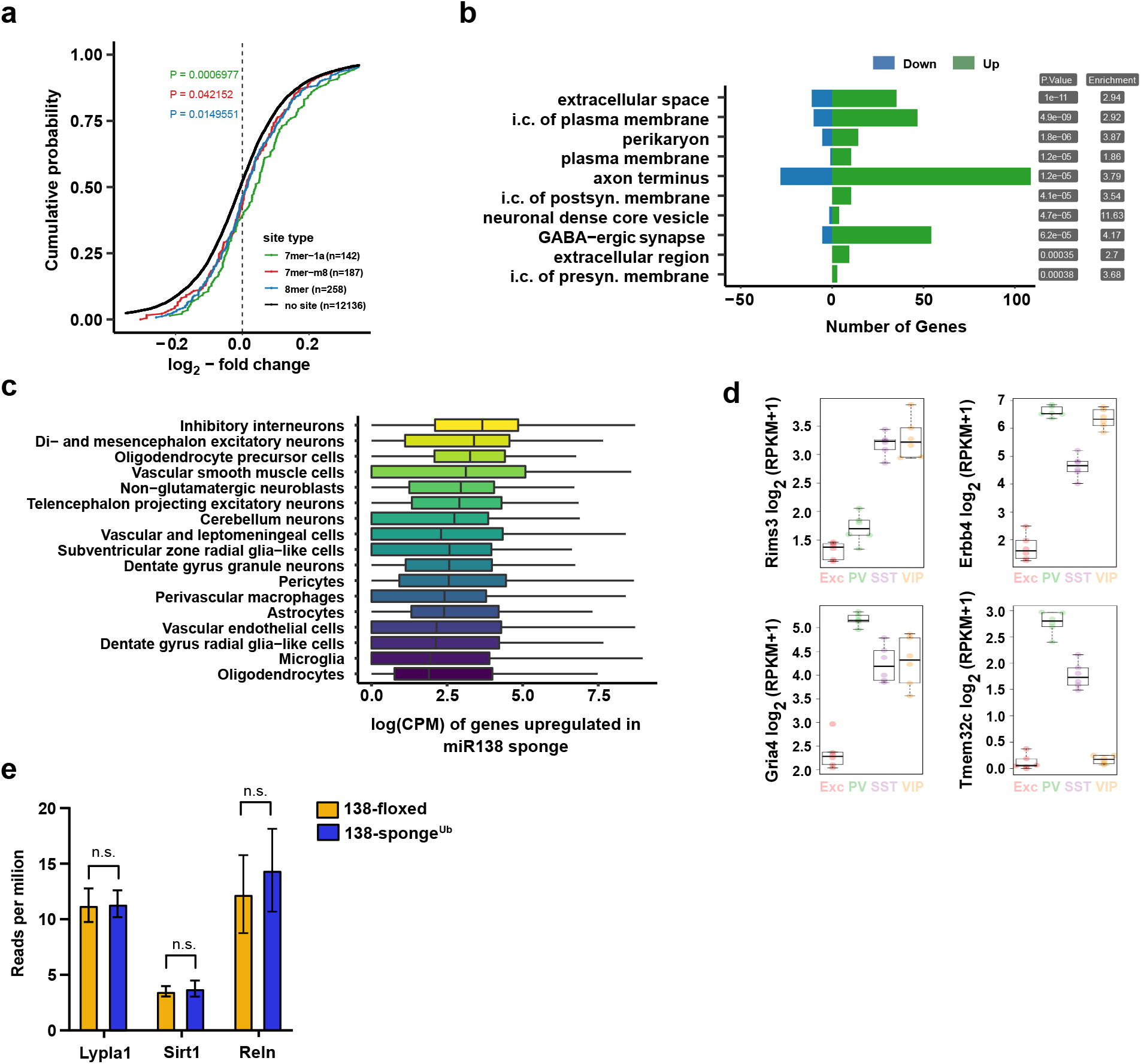
**(a)** Cumulative distribution plots of log_2_-fold expression changes (138-sponge^ub^/138-floxed) for genes either containing 7mer-1a (green), 7mer-m8 (red), 8mer (blue) or no (no site, black curve) predicted miR-138 binding sites. P-value is calculated compared to the no site population and indicated in the graph (KS-test). **(b)** Gene ontology (GO) term analysis for DEGs. Top ten enriched cellular component (CC) GO terms are shown. **(c)** Normalized expression levels, in single-cell clusters from Zeisel et al. (2018), of the genes upregulated in the miR-138 sponge. The genes are most abundantly expressed in inhibitory neurons. **(d)** Expression plots of selected miR-138 binding site containing transcripts in different neuronal subtypes (EXC: excitatory neurons; PV: parvalbumin+ interneurons; SST: somatostatin+ interneurons; VIP: Vasoactive intestinal peptide+ interneurons based on http://research-pub.gene.com/NeuronSubtypeTranscriptomes/. **(e)** Expression of validated miR-138-5p targets based on RNA-seq in 138-floxed and 138-sponge^Ub^ mice. n=3 mice. Lypla1 n.s. p=0.90; Sirt1 n.s. p=0.66; Reln n.s. p=0.50 (student’s two-tailed heteroscedastic t test). **Suppl. Figure 2 – source data:** this file contains the raw data on which the graphs in suppl. Fig. 2 are based.

**Supplementary Figure 3.**
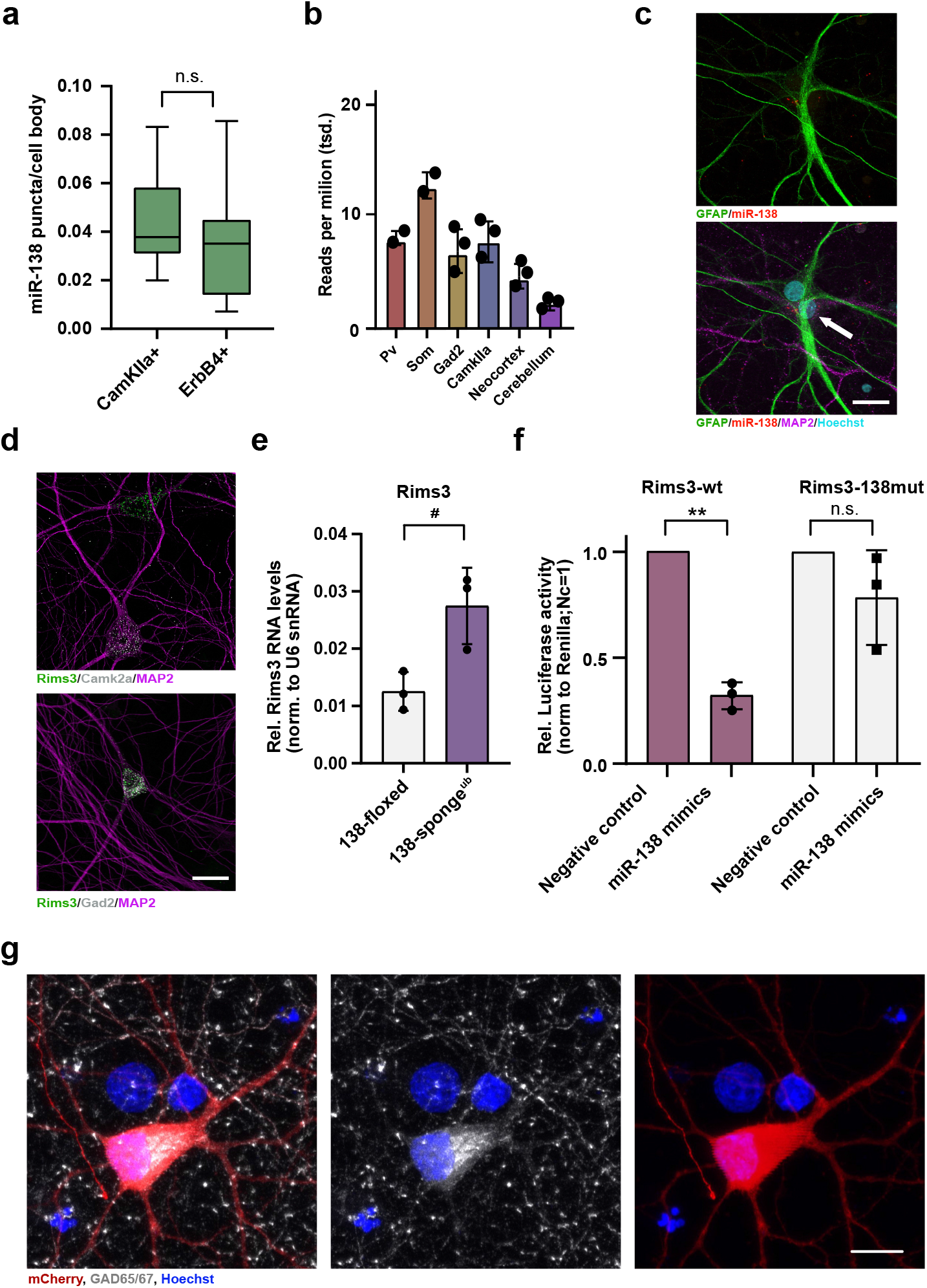
**(a)** Quantification of miR-138-5p smFISH in Camk2a- and Erbb4-positive neurons. Camk2a-positive n=10 cells; ErbB4-positive n=7 cells, p=0.48 (student’s two-tailed heteroscedastic t test). **(b)** Quantification of miR-138-5p levels in mouse brain tissue based on small RNA-seq data from (He et al., 2012) **(c)** Single-molecule FISH analysis of miR-138 (red) together with GFAP antibody stain (green) to label glia cells. Hoechst was used to counterstain nuclei. Arrow points to miR-138/MAP2 positive, GFAP-negative neuron adjacent to glial cell. Scale bar=20 µm. **(d)** Single-molecule FISH analysis of Rims3 mRNA (green) together with Camk2a (grey; left) or Gad2 (grey; right) mRNA to label glutamatergic and GABAergic neurons, respectively. Scale bar=20 µm. **(e)** qPCR analysis of Rims3 mRNA in total hippocampal RNA obtained from 138-floxed or 138-sponge^ub^ mice. U6 snRNA was used for normalization. n=3 mice; *p=0.04 (Student’s two-tailed heteroscedastic t test). **(f)** Relative luciferase activity in rat cortical neurons (DIV9-12) transfected with Rims3 3’UTR constructs with (138mut) or without (wt) a mutation in the miR-138 binding site, together with miR-138 or negative control mimics. Negative control mimic = 1. n=3 independent transfections, *p=0.002, n.s. p= 0.23 (Student’s two-tailed heteroscedastic t test). **(g)** Representative picture of primary rat hippocampal neurons infected with Dlx5/6-mCherry-138-sponge (red) and stained for GAD65/67 (grey) and Hoechst (blue). Scale bar = 10 μm. **Suppl. Figure 3 – source data:** this file contains the raw data on which the graphs in suppl. Fig. 3 are based.

**Supplementary Figure 4.**
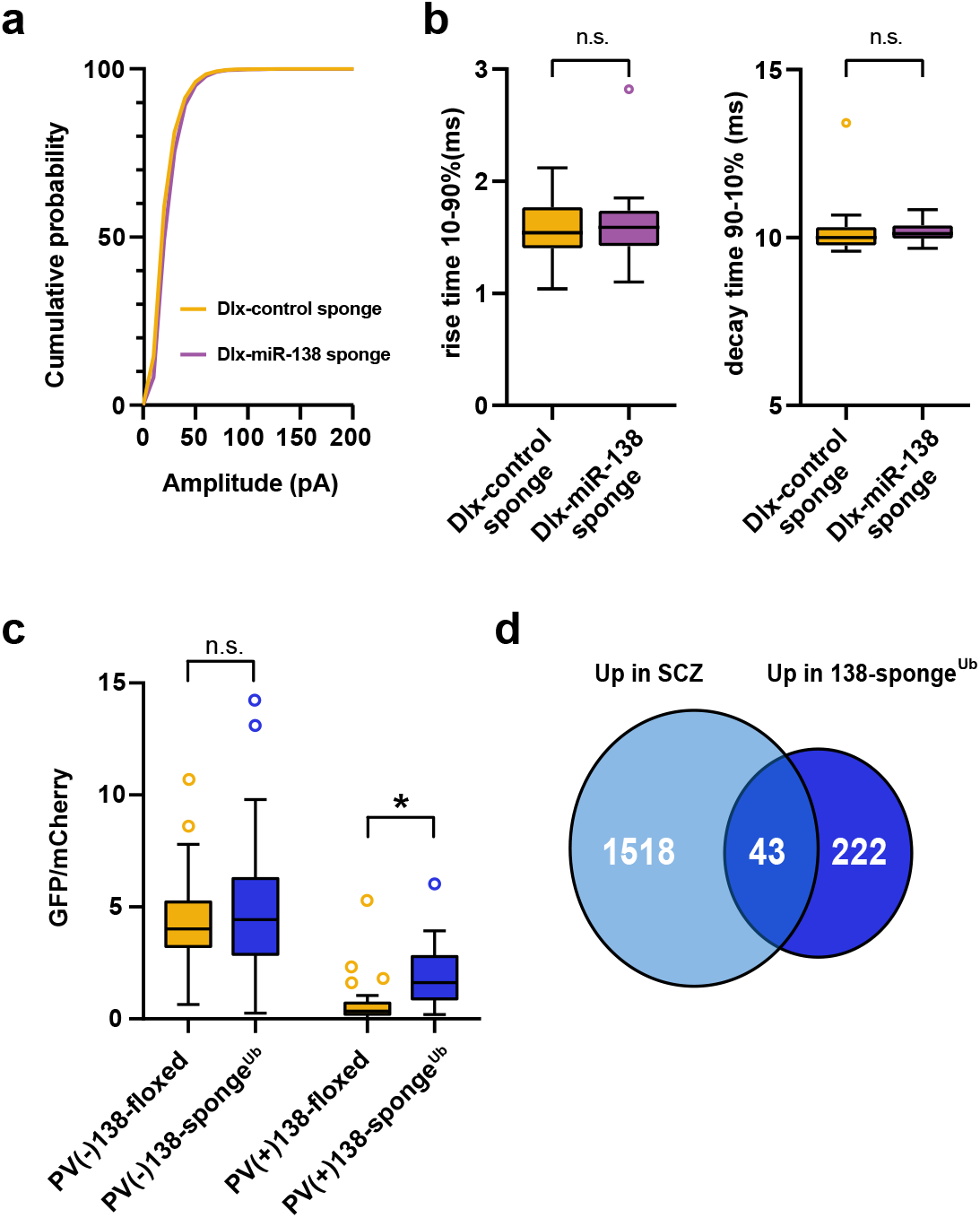
**a)** Cumulative distribution of mIPSC amplitude (p<0.0001; Kolmogorov-Smirnov test)**. b)** mIPSC rise (10-90%) and decay (90-10%) time. Left panel: mIPSC rise (10-90%) time (Dlx control sponge: range, from 1.0 to 2.1 ms; median, 1.5 ms; IQR, 0.4 ms. Dlx miR-138 sponge: range, from 1.1 to 2.8 ms; median, 1.6 ms; IQR, 0.3 ms; n.s. p=0.76 Mann Whitney test). Right panel: mIPSC decay (90-10%) time (Dlx control sponge: range, from 9.6 to 13.4 ms; median, 10.0 ms; IQR, 0.6 ms. Dlx miR-138 sponge: range, from 9.7 to 10.8 ms; median, 10.1 ms; IQR, 0.4 ms; n.s. p=0.3 Mann Whitney test). Dlx control sponge n=19 cells/2mice; Dlx miR-138 sponge n=19cells/2mice. **c)** Bar graphs of GFP/mCherry ratios from PV+ or PV- CA1 hippocampal neurons in 138-floxed or 138-sponge^Ub^ mice infected with a 138-pbds sensor construct; 138-floxed/PV+: n=32(2 mice); 138-floxed/PV-: n=40(2 mice); 138- sponge/PV+: n=32(3 mice); 138-sponge/PV-: n=42(3 mice); n.s. p=0.113;* p=3.6×10^-6^ (Kolmogorov-Smirnov-Test). d) Venn diagram describing the overlap between transcripts upregulated in SCZ patients (Gandal et al., 2018) and miR-138 sponge mice (Fig. 2a). Fold enrichment = 1.43; p=0.00859 (Fisher test). **Suppl. Figure 4 – source data:** this file contains the raw data on which the graphs in suppl. Fig. 4 are based.

**Supplementary Figure 5.**
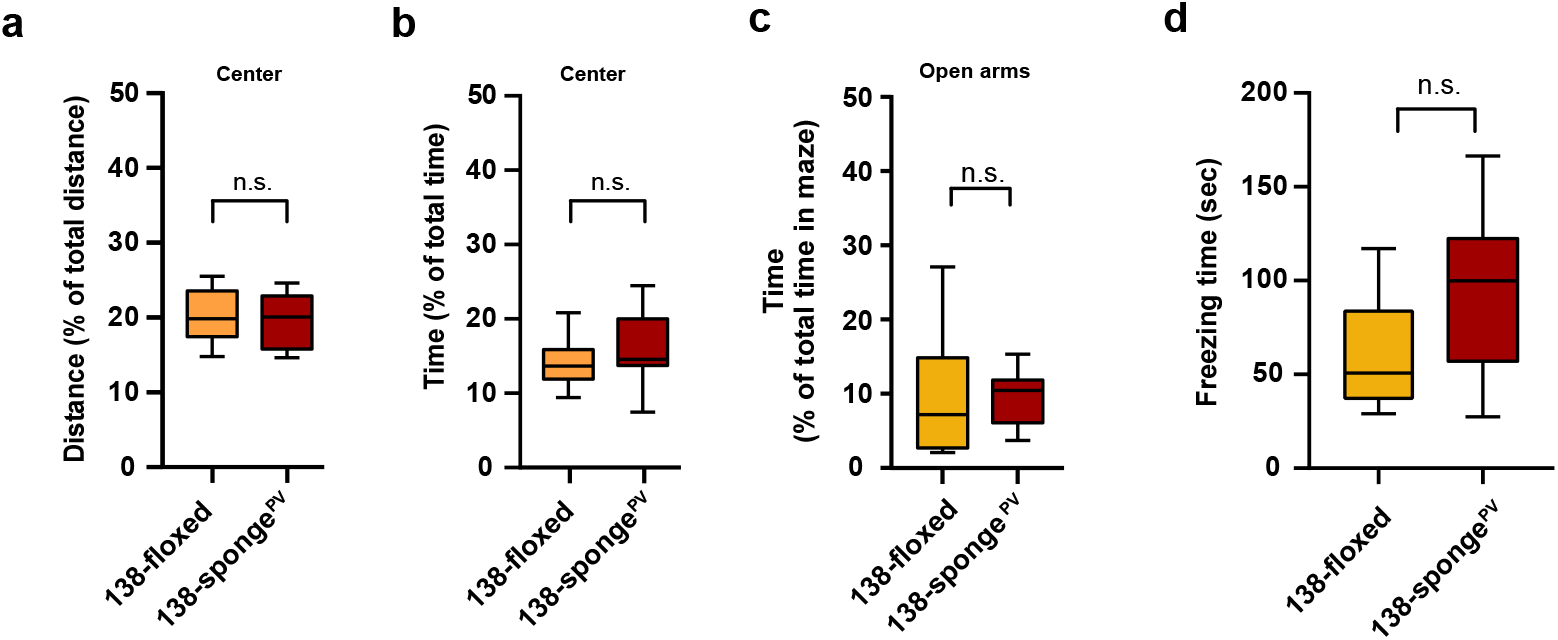
**(a-b)** Percentage of total distance travelled (a)/time spent (b) in the center of an open field arena during 30 min exploration by mice of the indicated genotypes; 138-floxed n=10 mice; 138-sponge^PV^ n=10 mice; (a) p=0.74 (b) p=0.28 (Student’s two-tailed heteroscedastic t-test). **(c)** Percentage of total time spent in the open arms of an elevated plus maze (EPM) during 5 min exploration by mice of the indicated genotypes; 138-floxed n=10 mice; 138-sponge^PV^ n=10 mice; p=0.53 (Mann-Whitney test). **(d)** Time (s) mice spent freezing 24 h after the foot shock was administrated; 138-floxed n=9 mice; 138-sponge^PV^ n=9 mice; p=0.075 (Student’s two-tailed heteroscedastic t-test). **Suppl. Figure 5 – source data:** this file contains the raw data on which the graphs in suppl. Fig. 5 are based.

**Supplementary Figure 6.**
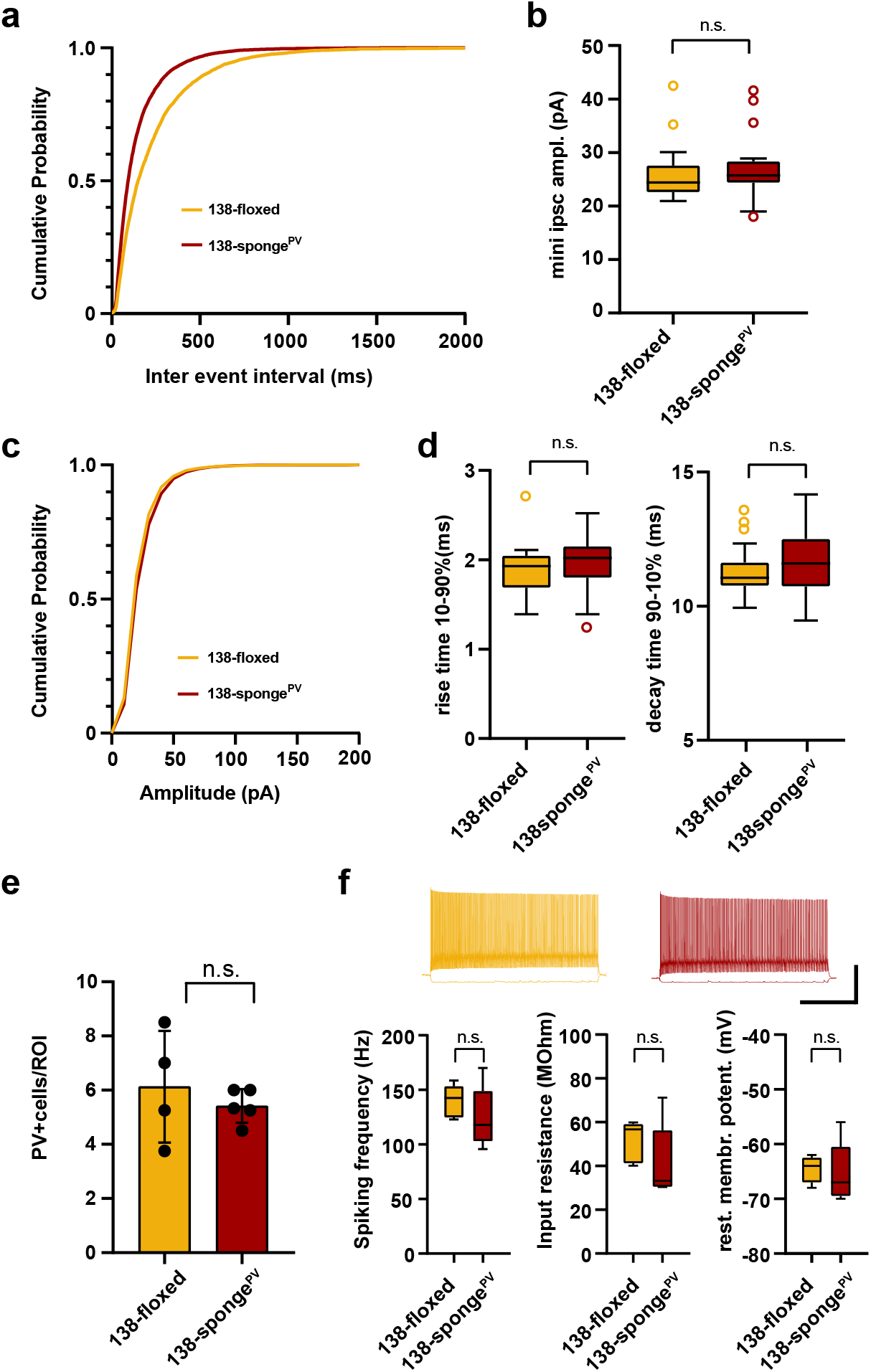
**(a-d)** mIPSC in CA1 pyramidal neurons. **(a)** Cumulative distribution of mIPSCs frequency. (p<0.0001; Kolmogorov-Smirnov test) **(b)** mIPSC amplitude (138-floxed: range, from 20.9 to 42.5 pA; median, 24.4 pA; IQR, 5.0 pA. 138-sponge^PV^: range, from 18.0 to 41.6 pA; median, 25.7 pA; IQR, 4.0 pA; n.s. p=0.22 Mann Whitney test). **(c)** cumulative distribution of mIPSCs amplitude (p<0.0001 Kolmogorov-Smirnov test). 138-floxed n=22 cells/5mice; 138-sponge^ub^ n=23cells/5mice. **(d)** mIPSC rise (10-90%) and decay (90-10%) time. Left panel: mIPSC rise (10-90%) time (138- floxed: range, from 1.4 to 2.7 ms; median, 1.9 ms; IQR, 0.4 ms. 138-sponge^PV^: range, from 1.3 to 2.5 ms; median, 2.0 ms; IQR, 0.4 ms; n.s. p=0.28 Mann Whitney test). Right panel: mIPSC decay (90-10%) time (138-floxed: range, from 9.9 to 13.6 ms; median, 11.1 ms; IQR, 0.9 ms. 138-sponge^PV^: range, from 9.5 to 14.2 ms; median, 11.6 ms; IQR, 1.8 ms; n.s. p=0.20 Mann Whitney test). 138-floxed n=22 cells/5mice; 138-sponge^ub^ n=23cells/5mice. **(e)** Density of PV+ interneurons in CA1 hippocampus of mice with the indicated genotype. Values are expressed relative to a defined region of interest (ROI). 138-floxed: n=16 ROIs/4 mice; 138-sponge^PV^: n=18 ROIs/5 mice; data represents the average per mouse ± s.d.; n.s. p=0.8049 (Mann-Whitney test). **(f)** Properties of fast-spiking interneurons. Upper panel: example traces, 138-floxed in orange, 138-songe^PV^ in red, scale bar: 50 mV, 500 ms. Lower panel left: spiking frequency (138-floxed: range, from 123 to 159 Hz; median, 143 Hz; IQR, 29 Hz. 138-sponge^PV^: range, from 96 to 170 Hz; median, 118 Hz; IQR, 46 Hz; n.s. p=0.31 Student’s two-tailed heteroscedastic t test). Lower panel middle: input resistance (138-floxed: range, from 40 to 60 MΩ; median, 57 MΩ; IQR, 18 MΩ. 138-sponge^PV^: range, from 30 to 71 MΩ; median, 33 MΩ; IQR, 26 MΩ; n.s. p=0.22 Mann Whitney test). Lower panel right: resting membrane potential (138-floxed: range, from -68 mV to -62 mV; median, -64 mV; IQR, 4.5 mV. 138-sponge^PV^: range, from -70 to -56 mV; median, -67 mV; IQR, 9 mV; n.s. p=0.78 Student’s two-tailed heteroscedastic t test). 138-floxed n=5 cells/3mice; 138-sponge^ub^ n=5cells/5mice **Suppl. Figure 6 – source data:** this file contains the raw data on which the graphs in suppl. Fig. 6 are based.

